# Rich dynamics and functional organization on topographically designed neuronal networks *in vitro*

**DOI:** 10.1101/2022.09.28.509646

**Authors:** Marc Montalà-Flaquer, Clara F. López-León, Daniel Tornero, Tanguy Fardet, Pascal Monceau, Samuel Bottani, Jordi Soriano

## Abstract

Neuronal cultures are a prominent experimental tool to understand complex functional organization in neuronal assemblies. However, neurons grown on flat surfaces exhibit a strongly coherent bursting behavior with limited functionality. To approach the functional richness of naturally formed neuronal circuits, here we studied neuronal networks grown on polydimethylsiloxane (PDMS) topographical patterns shaped as either parallel tracks or square valleys. We followed the evolution of spontaneous activity in these cultures along 20 days *in vitro* using fluorescence calcium imaging. The networks were characterized by rich spatiotemporal activity patterns that comprised from small regions of the culture to its whole extent. Effective connectivity analysis revealed the emergence of spatially compact functional modules that were associated to both the underpinned topographical features and predominant spatiotemporal activity fronts. Our results show the capacity of spatial constraints to mold activity and functional organization, bringing new opportunities to comprehend the structure-function relationship in living neuronal circuits.

## INTRODUCTION

A fascinating yet intriguing property of living neuronal circuits is their capacity to exhibit a rich repertoire of activity patterns and functional states from a relatively hardwired structural architecture (Blankenship and Feller, 2010; Suárez et al., 2020). This property is most prominent in the human brain, enabling the realization of precise and fast-changing tasks with precision, from motor action to memory and cognition (Finc et al., 2020; Park and Friston, 2013), and that reveals the existence of intrinsic mechanisms and network traits for a swift dynamic reconfiguration of neural circuits. An established consensus is that modular and hierarchically modular network organization (Meunier et al., 2010) are fundamental hallmarks for the coexistence of diverse dynamic scenarios, allowing for both specialized computation at the scale of a module (functional segregation) and whole-network information exchange (functional integration) (Deco et al., 2015; Sporns, 2013) with balanced wiring-efficiency cost (Bullmore and Sporns, 2012).

Modularity and integration-segregation balance are important actors in the functioning of neuronal circuits and play a key role in their robustness and flexibility (Finc et al., 2020). The sheer size of the brain and the intrinsic difficulty to monitor neuronal-level dynamics in detail, however, have fostered the development of *in vitro* preparations in which complex behavior at the mesoscale can be investigated (Aebersold et al., 2016; Millet and Gillette, 2012; Wheeler and Brewer, 2010). Culturing neurons in a controlled environment allows not only for an easy accessibility and manipulation of ∼100 − 1000 neurons but also for the design of true ‘structure-to-function’ laboratories to investigate the relation between physical wiring and emerging complex behavior (Bonifazi et al., 2013; Marconi et al., 2012; Poli et al., 2015).

Different experimental studies have pointed out the advantage of spatial constraints, connectivity guidance and modular designs (Neto et al., 2016) to tune neuronal culture functionality (Forró et al., 2018) and dynamics (Bisio et al., 2014; Park et al., 2021), or to facilitate the coexistence of segregated and integrated states (Yamamoto et al., 2018). The advent of three-dimensional cultures have also paved the way towards elaborate ‘brain-on-chip’ devices that aim at reproducing and modeling *in vitro* the building blocks of complex brain behavior (Pas, 2018; Zhuang et al., 2018). Despite the importance of these neuroengineering efforts, it has been shown that an initially homogeneous distribution of neurons undergo significant reorganization that shape complex network features such as small-worldness and rich-club topology (Downes et al., 2012; Schroeter et al., 2015), whereas cultures exhibiting mild fluctuations in the spatial distribution of neurons in combination with activity-depend mechanisms evolve to exhibit modularity traits and balanced local-to-global connectivity (Okujeni and Egert, 2019; Tibau et al., 2020).

Although the above studies show that self-organization in neuronal circuits suffice to imprint rich functional traits, an aspect that remains unexplored is whether these traits can be accelerated or strengthened by incorporating coarse spatial constraints that break the isotropy of the substrate in which neurons grow. To advance in this quest, here we used mesoscopic neuronal cultures 6 mm in diameter grown on PDMS topographical substrates that contained elevations shaped as either parallel tracks or squares. Using effective connectivity and complex networks analyses we show that the underlying topography alters the way in which neurons develop and interconnect, shaping a rich repertoire of activity patterns and functional traits — most notably modularity— that contrast with the strongly coherent behavior and weak modularity of standard cultures grown on a flat surface. Our work provides compelling evidence that spatial constraints and structural features mold whole-network spontaneous activity and functional organization, opening new avenues for understanding the structure-function relationship in neuronal assemblies.

## RESULTS

### PDMS topographical molds enrich spontaneous activity in primary neuronal cultures

We used printed board technology to generate a master mold formed by copper motifs 70 µm high deposited on a fiberglass substrate (Figures 1A and S1). As described in Transparent Methods, the motifs were designed using computer-aided design software in combination with printed circuit board technology an included two main configurations, namely parallel copper tracks (termed *tracks*) and randomly positioned square copper blocks (*squares*). As sketched in Figure 1A, using copper tracks as example, the printed circuit board shaped a relief over which PDMS was poured and cured, giving rise to a topographical design that was the negative of the original mold. PDMS was then cut out as discs 6 mm in diameter that were firmly attached to a glass coverslip, and primary neuronal cultures from rat embryos were grown on the PDMS surface in a homogeneous manner. Cultures were later transduced with the genetically encoded calcium indicator GCaMP6s using adeno-associated viruses (AAVs) and spontaneous activity was monitored using calcium fluorescence imaging. Measurements on the same culture extended from day *in vitro* (DIV) 7, in which fluorescence signal was sufficiently strong for reliable analysis, to DIV 18, in which neurons started to degrade or detach from the PDMS surface.

**Figure 1.**
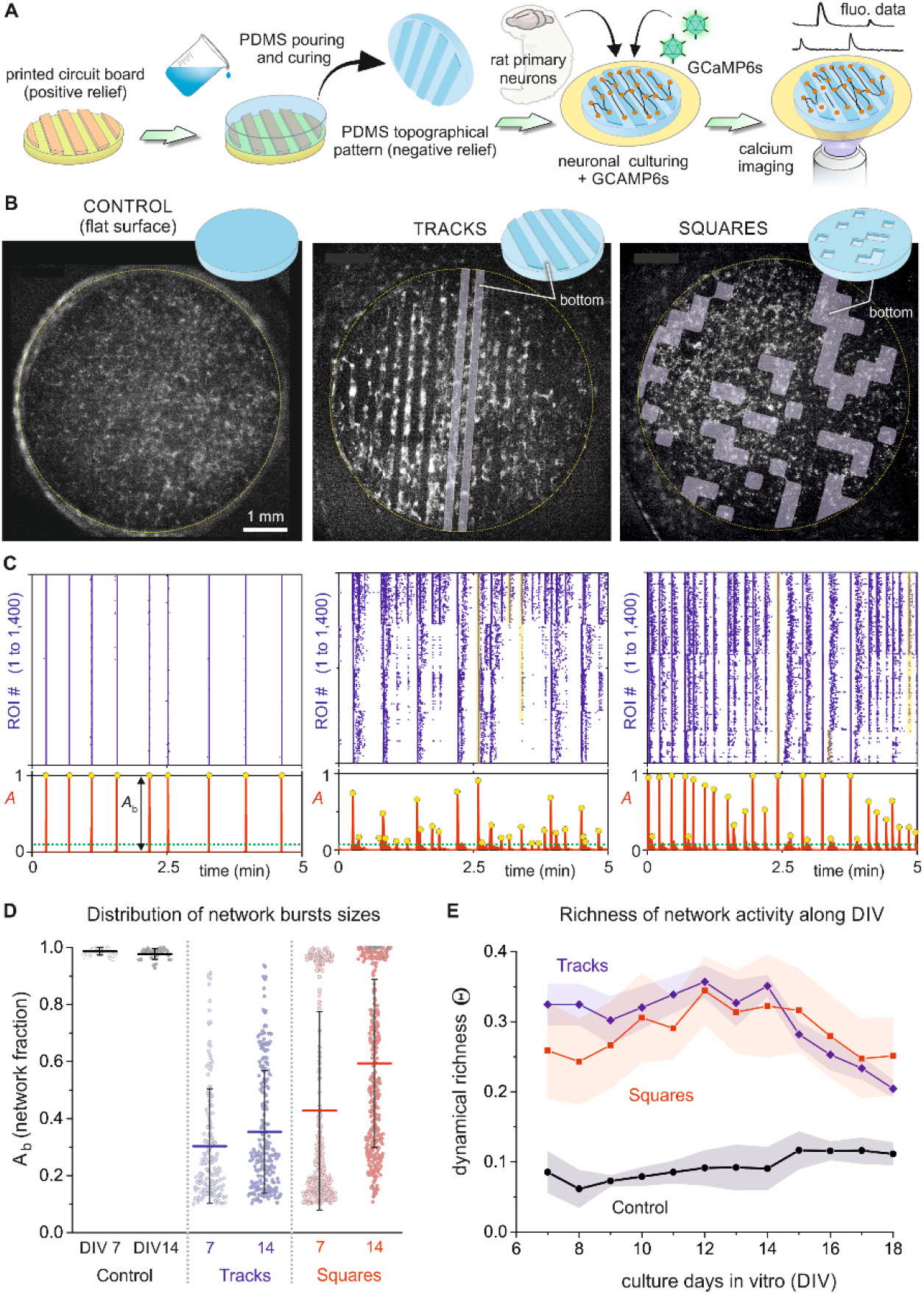
Spontaneous activity of neuronal cultures grown on PDMS topographical molds. (A) Sketch of the experimental setup and procedure. A printed board circuit with a topographical relief 70 µm high (orange) was used as a master mold to pour and cure PDMS on it, leading to a design (blue) that is the negative relief of the original mold. Neurons were cultured on it in combination with GCaMP6s, delivered through adeno-associated viruses (AAVs). Spontaneous neuronal activity was then monitored through calcium fluorescence imaging. (B) Illustrative fluorescence images of the three studied topographical designs, namely a flat PDMS surface that serves as control (left), parallel tracks (center) and randomly positioned square valleys (right). All cultures were 6 mm in diameter and were recorded at DIV 14. Bright spots on the fluorescence images reveal active neurons. (C) Corresponding raster plots (top) and population activity *A* (bottom). Events encompassing more than 10% of the monitored Regions of Interest (ROIS) (green lines) were considered significant (yellow dots) and shaped ‘network bursts’ of size *A*_*b*_. Raster plots were ordered by ROIs similarity to highlight groups of coordinated activity. (D) Distribution of bursting sizes *A*_*b*_ for the three configurations and comparing young (day *in vitro*, DIV, 7) and mature (DIV 14) cultures. Each box plot contains all experimental repetitions for a given condition. (E) Dynamical richness ⨸ along development. ⨸ portrays the variability in the raster plots, which is much higher in topographical cultures, particularly in the range DIV 7-14. Data in panels (D) and (E) are based on n=5 cultures for flat PDMS, 5 for tracks, and 6 for squares.

Figure 1B provides illustrative fluorescence images of the prepared neuronal cultures. We included in our study control cultures plated on a flat PDMS surface to better assess the impact of topography on neuronal network behavior. To quantify the collective behavior of the prepared cultures, we recorded spontaneous activity in each configuration for 30 min, to next extract the fluorescence traces in small Regions of Interest (ROIs) that contained 5-10 neurons each and that covered uniformly the area of the culture (Figure S2), giving rise to about 1,400 ROIs.

The recorded fluorescence data was analyzed to extract the activation time of each ROI, and data represented in the form of raster plots. Figure 1C shows 5 min of representative data for the cultures depicted in Figure 1B. The corresponding calcium imaging recordings are provided in Supplementary Videos SV1 to SV3. Activity in control cultures was characterized by episodes of highly coherent behavior in which all ROIs activated together in a short time window of ∼200 ms (*network bursts*) or remained practically silent. *Tracks* and *squares* configurations, by contrast, showed a much richer dynamic repertoire, in which network bursts of different sizes coexisted (yellow bands in Figure 1C). Network bursts extended longer periods of time (on the order of seconds) for these topographical configurations, and sporadic activity outside bursts was also more abundant.

The rich variety of network burst sizes was reflected in the population activity *A*, which counts the fraction of ROIs that coactivate together (Figure 1C, bottom panels). Networks bursts whose size were above background activity (typically 10% of *A*) were considered significant and denoted *A*_*b*_. While all events exhibited sizes *A*_*b*_ = 1 for controls, the event sizes for the topographical designs richly varied from *A*_*b*_ ≳ 0.1 to *A*_*b*_ ≃ 1.

A comparison of the distribution of bursting sizes in the different configurations is provided in the boxplots of Figure 1D. Data incorporate different experimental repetitions for the same configuration and compare young (DIV 7) and mature (DIV 14) cultures. For young cultures, while controls produced a narrow distribution with ⟨*A*_*b*_ ⟩ ≃ 1, tracks and squares were clearly peaked toward small values of *A*_*b*_, with ⟨*A*_*b*_ ⟩ ≃ 0.3 and 0.35, respectively. Upon maturation, the distribution of bursting sizes remained practically the same for controls, indicating that these cultures activate in a coherent manner in all its lifespan. For tracks and squares, the distributions shifted towards higher *A*_*b*_ values, with ⟨*A*_*b*_ ⟩ ≃ 0.4 and 0.6, respectively. We argue that this increase in busting sizes upon maturation is associated with an overall stronger interconnectivity in the network, which reduces the impact of topography on dynamics.

The variety in activity patterns, which is reflected both in the structure of the raster plots and the distribution of *A*_*b*_ values, can be quantified through a single parameter termed *dynamical richness* Θ (Yamamoto et al., 2018) which varies between Θ = 0 for perfectly coherent or random activity and Θ = 1 for maximally patterned activity, *i*.*e*., with all possible neuronal coactivation patterns present, from few neurons to the entire network. Figure 1E shows the results for the evolution of Θ as a function of DIV for the three configurations, averaged out among different repetitions. While Θ ≲ 0.1 for controls, with small changes along development, Θ exhibits at short DIV a much larger Θ ≃ 0.33 and Θ ≃ 0.25 for tracks and squares, respectively, remaining high for about a week until it decreases after DIV 14, as overall activity in the topographical designs becomes globally more coherent.

### Topographical cultures give rise to a rich repertoire of spatiotemporal activity patterns

The burst size *A*_*b*_ captures the fraction of the network that activates coherently in a short time window but does not inform about the spatiotemporal structure of a burst. To explore this aspect, Figure 2A provides image plots of representative bursts’ evolution across the culture for the different configurations. For tracks and squares, we included data for young (DIV 7) and mature (DIV 14) cultures since they changed in dynamic behavior along maturation. This was not the case for control cultures, which exhibited whole-network bursts in all their evolution, and therefore only data at DIV 7 is shown. In the image plots we used a black-yellow color scheme to portray the activation time of each ROI, and marked the initiation point of the burst spatiotemporal front as a white circle. Regions of the culture with no activity are shown in grey.

**Figure 2.**
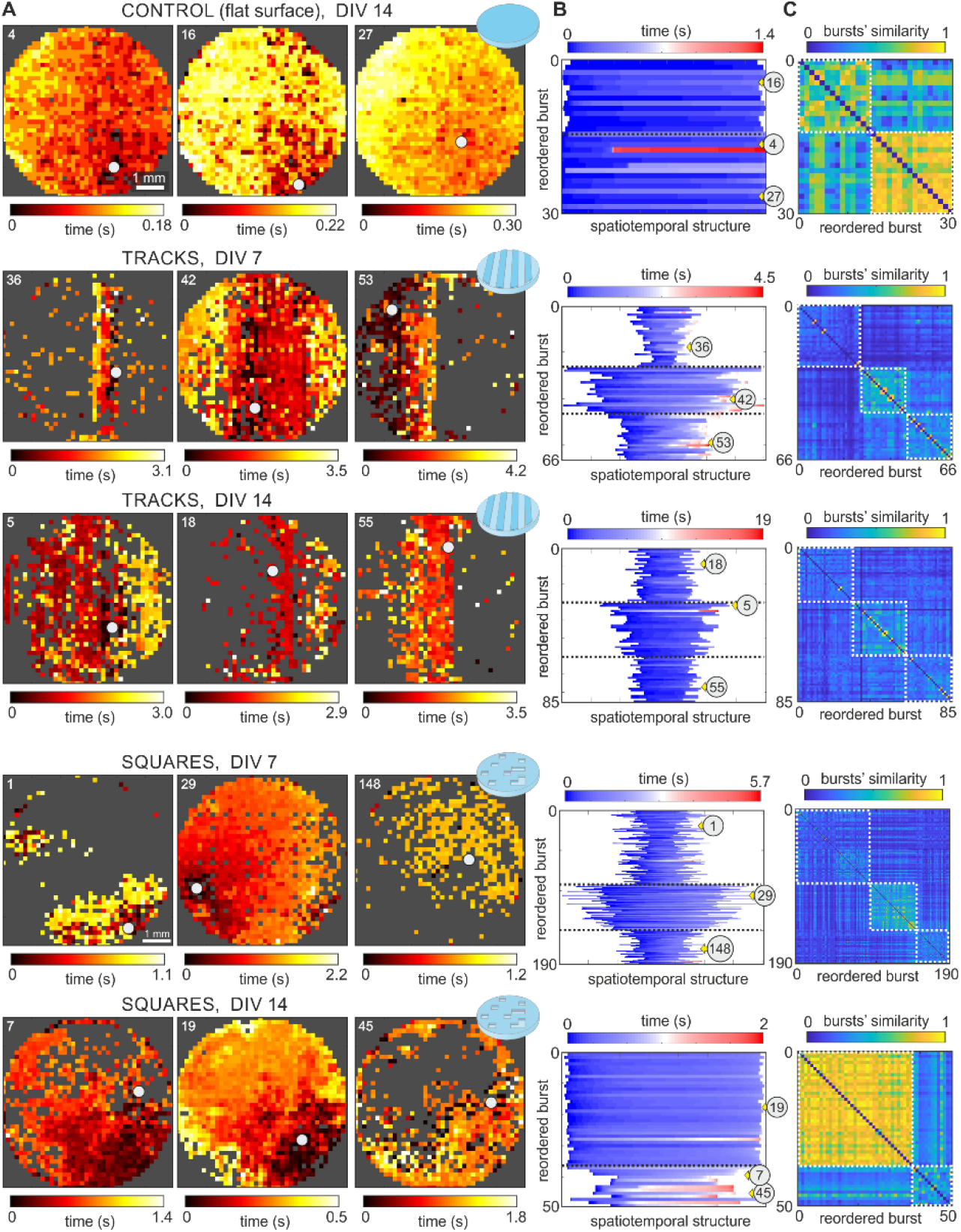
Repertoire of spatiotemporal patterns in control and topographical cultures. (A) Representative examples of spatiotemporal activity fronts for controls, tracks and squares configurations. For the two latter, both young and mature developmental stages are shown for the same culture. Each colored dot in the image plots is an active ROI, with the color coded according to the time of activation (from black to yellow). Grey regions indicate absence of active ROIs. Activity in controls always comprises the entire network and propagates fast (*Δt* ≃ 0.2 − 0.3 s). For lines and squares, activity switches between few sections of the culture or its whole extent, and much slowly (*Δt* ≃ 1 − 3 s). The number on the top-left corner of each panel identifies the burst, while the big white dot signals the origin of activity. For the tracks configuration, the image plots are drawn for the topographical tracks to coincide with the vertical direction. For the squares configuration, the motifs are aligned with the image borders. (B) Classification of burst sizes and propagation. Each band corresponds to a burst, which are ordered according to their similarity. The width of a band indicates the number of ROIs involved in the burst, while its color indicates the propagation time. Bands with similar color scheme portray bursts with akin spatiotemporal structure. The numbers within a grey circle indicate the position of the bursts represented in (A). The doted black lines separate the different groups of bursts. (C) Detailed classification in the form of similarity matrix and following the same organization as in (B). The brighter the color, the higher the similarity among bursts. White dashed boxes identify the groups of similar bursts.

For controls, activity encompassed the entire culture in all bursts and propagated as a quasi-circular front, which is reflected by a progressive change in color (from black to yellow) for ROIs gradually further from the origin of activity. Activity propagation was fast, with the bursts crossing the 6 mm diameter of the culture in about 0.2 − 0.3 s. Bursts also started approximately in the same location. Thus, the whole-network activation and the similar location of burst initiation shaped altogether a very rigid system. In the tracks configuration, by contrast, different sizes and propagation schemes were present in combination with a richer variability in initiation points. Indeed, at DIV 7, some bursts encompassed just a couple of tracks (front #36), the entire culture (#42) or half of it (#53). Propagating fronts required about 3 − 4 s to propagate over the culture, *i*.*e*., bursts were an order of magnitude slower than in controls. As the culture matured, the bursts maintained this variability in sizes and initiation points, although there was a tendency for the sizes to encompass larger areas. Bursts extending only one or two tracks were rare in these mature networks. For squares, we observed that young cultures exhibited a rich variability in burst structures, which encompassed either specific regions of the culture (bursts #1 and #148) or its entireness (#29). Burst propagation took about 1 − 2 s to cross the system, *i*.*e*., in between controls and tracks. The most prominent characteristic of the squares configuration, however, is that for mature cultures at DIV 14 there was a tendency for the bursts to cover large areas. Fragmented activity was rare (bursts #7 and 45) and most of bursts filled the entire culture (#19).

In Figure 2B we provide a diagram that compares in a compact manner the spatiotemporal structure of all bursts for each configuration and day *in vitro*. As explained in Transparent Methods, each color band in the diagram represents a burst. The width of the band indicates the number of participating neurons in each burst, while the color scale itself portrays the spatiotemporal evolution. Conceptually, those bursts that propagate similarly share the same color structure. The bursts are ordered in the y-axis according to a similarity analysis that identifies groups of akin bursts. Similarity was based on Pearson’s correlation among all pairs of bursts in combination with community detection (Figure 2C, white boxes). For Figure 2B, the groups of similar bursts are separated by a black line, and the gray discs with a number show the id of the bursts portrayed in Figure 2A. For controls, all bursts practically comprised the whole network and therefore they fill the width of the diagram. Additionally, most of the color bands evolve from dark blue to clear blue, indicating a similar activity propagation across the culture. For tracks at DIV 7 three distinct groups appeared in the diagram and were associated to activity extending a couple of tracks (top group), most of the culture (central group) or half of it (bottom group). The color scheme of the bursts was richer than in controls, indicating that spatiotemporal propagation was more varied. These three distinct groups were preserved upon maturation, although the groups were more similar among themselves and color schemes were more uniform. A similar trend was observed in the squares configuration. Three distinct groups of bursts were clear at DIV 7, which correspond to typically small yet compact areas of the culture (top), quasi full-culture activations (center), and small activations in scattered areas (bottom). These groups practically vanished at DIV 14 as most of the bursting events comprised the entire culture.

### Immunostaining reveals connectivity traits induced by the PMDS topography

To understand the origin of the rich repertoire of activity patterns, we carried out an immunohistochemical analysis on the tracks and squares configurations. As shown in Figure 3, we were interested in identifying neuronal processes (green), astrocytes (red) and cell nuclei (blue). For tracks (Figure 3A), confocal images covering a field of view on the order of mm (‘overview’, left) revealed that neuronal processes extended preferentially along the direction of the tracks, both at the top and at the bottom of the PDMS relief, and that connectivity in the transverse direction was by comparison very minor. Immunostaining also revealed that neuronal processes often tended to follow the edges of the relief, particularly at the top of the design (Figure 3A, detail, white arrowheads). Thus, topography provided guidance to connections, which were funneled along the tracks and shaped a highly anisotropic connectivity. A detail of the images (Figure 3A, right) allows to clearly visualize the difference in connectivity along and across tracks. This difference provided the seed for shaping track-oriented, weakly coupled microcircuits that ultimately rendered rich spatiotemporal patterns. The detail immunostaining images also reveal the abundant and uniform distribution of astrocytes in the cultures, which contrasts with the aggregation of cell nuclei (Figure 3A, bottom), a trait that could also help to enrich connectivity microcircuits and varied emerging dynamics.

**Figure 3.**
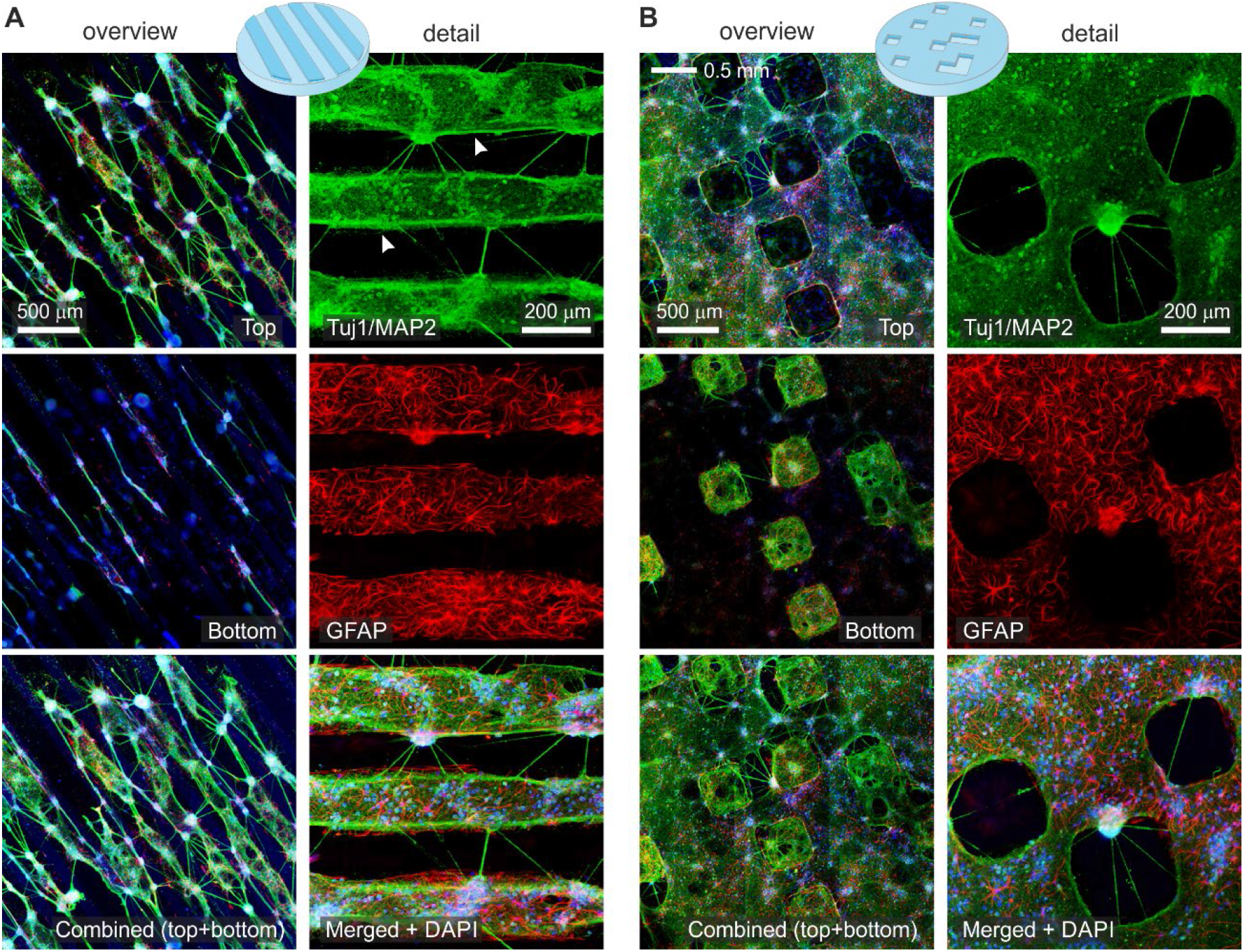
Anisotropic connectivity in PDMS topographical cultures revealed by immunostaining. (A) Representative immunohistochemical images of neurons grown on PDMS topographical tracks at DIV 14, providing a broad overview (left) and a detail (right). For the overview, images show neuronal processes in green, astrocytes in red and cell nuclei in blue, with focus at the top part of the topography, the bottom part, and a combination of them. For the detail, images correspond to the top part of the design only and depict neuronal process (green), astrocytes (red), and the combination of these channels with cell nuclei (blue). Neuronal connections are more abundant in the direction of the tracks than transverse to them. White arrows mark neuronal processes that follow the edge of the topographical design. (B) Corresponding images for the squares configuration at DIV 14. Connectivity is abundant at the bottom part of the square designs, shaping small microcircuits by themselves that interconnect with the top part.

The immunohistochemical analysis for the squares configuration is provided in Figure 3B, both at the mm scale (left) and in the detail (right). For this configuration, the relief favored the formation of strongly connected islands at the bottom part of the relief which, in turn, connected with other islands or with the top part of the PDMS. Astrocytes and neurons were also abundant, although the latter were more homogenously distributed as compared to tracks. Thus, for squares, the presence of spontaneous activity as ‘patches’ in Figure 2A (squares, DIV7) is most likely due to the activation of relatively isolated groups of neurons either at the top or at the bottom of the topographical relief. The interaction among the top and bottom parts, however, is complex since the entire network sporadically activated in a coherent manner. We note that, as in tracks, the relief facilitates the formation of interacting distinctive microcircuits that shape a rich variety of spontaneous activity patterns. These microcircuits are not stable in time, but rather continuously evolve. Indeed, the observation that activity at DIV 14 mostly encompasses the entire network indicates that there is global tendency for the microcircuits to gradually blend together and cast a much more uniform overall connectivity that translates into whole-network bursting events.

### PDMS topography impacts on burst initiation and velocity of burst propagation

Figure 4A provides the spatial distribution of burst initiation for the configurations shown in Figure 2 at two stages of culture maturation (DIV 7 and DIV14). In the panels, the black dots represent the spatial location of each observed burst while the blue-yellow colormap shows the corresponding probability distribution function of burst initiation. The degree of activity focalization, *i*.*e*., the tendency for spontaneous activity to initiate in the same location, is quantified through the Gini coefficient λ, which is 0 for a spatially equiprobable initiation, and 1 for a point-concentrated initiation occurrence. For controls (left), most of the bursts at DIV 7 started in the same neighborhood, leading to highly focalized distribution function (yellow spot) with λ ≃ 0.57. This focalization was maintained at DIV 14 (λ ≃ 0.51), although the location of the most probable initiation points varied due to global connectivity changes during maturation. By contrast, activity initiation for the tracks configuration was substantially more extended at DIV 7 (λ ≃ 0.20), a trait that was maintained upon maturation (λ ≃ 0.16 at DIV 14). Hence, topographical tracks not only help shaping connectivity anisotropies that enriched spontaneous activity but that these anisotropies were maintained upon maturation. For squares, initiation was spatially extended at DIV 7 (λ ≃ 0.28) but became focalized upon maturation (λ ≃ 0.72 at DIV 14). This focalization is consistent with the observed whole-network bursting and the gradual loss of connectivity anisotropies upon maturation.

**Figure 4.**
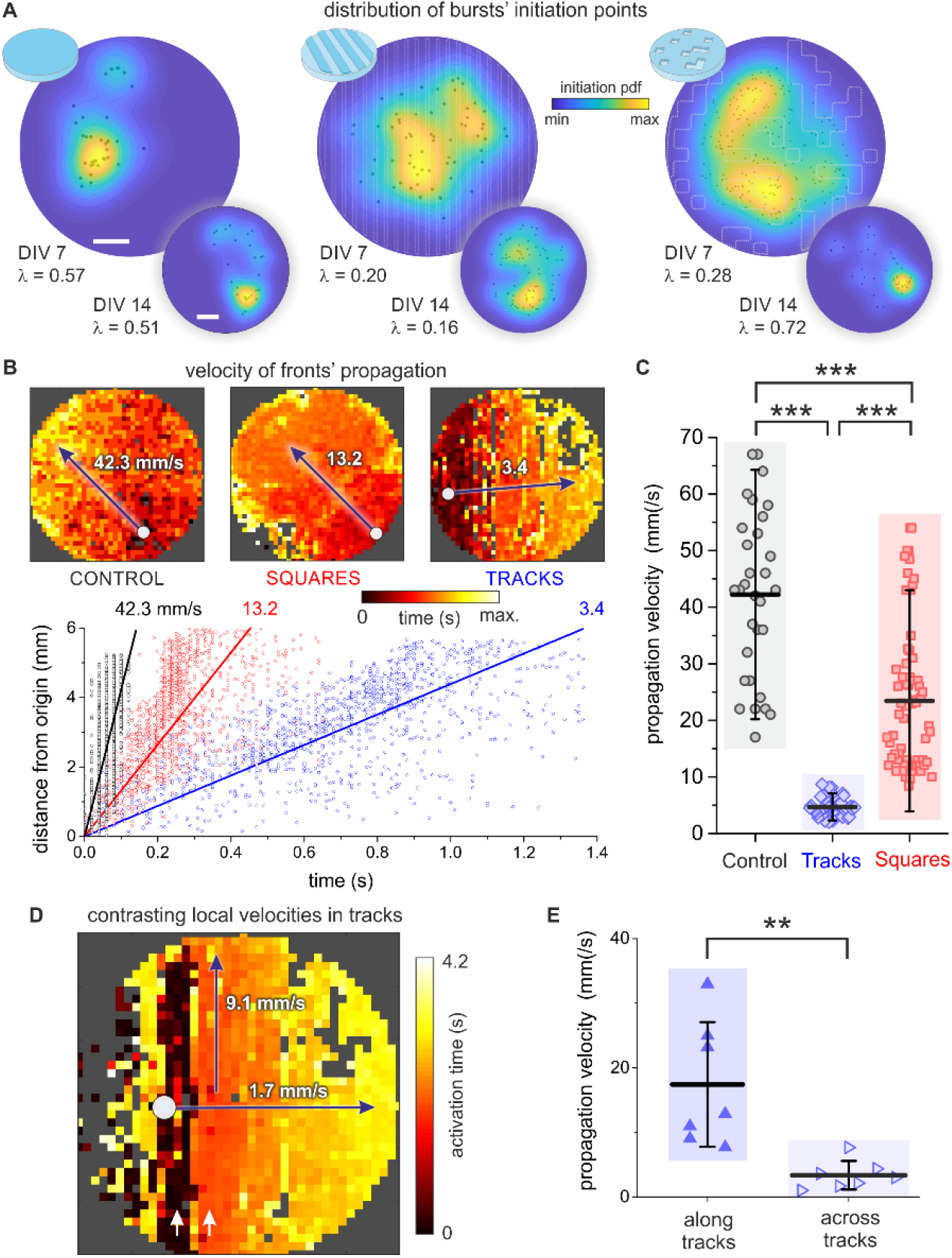
Burst initiation and velocity of propagation bursts in PDMS topographical cultures. (A) Spatial distribution of burst initiation points (black dots) and probability density functions of burst initiation (PDF, blue-yellow colormap) for the PDMS configurations shown in Figure 2 and comparing two days *in vitro*, DIV 7 and DIV 14. *λ* is the Gini coefficient and indicates the degree of focalization of burst initiation. Scale bars are 1 mm. (B) Top, representative network bursts that encompass the entire network and whose propagation is compatible with a circular or flat front. The white dot marks the origin of activity. Bottom, corresponding determination of velocity propagation as linear fits, where the Euclidean distance of each ROI to the origin of activity is plotted as a function of its activation time. All linear fits have Pearson’s correlation coefficients *r* ≳ 0.95. (C) Box plots of propagation velocities for the three configurations, showing that fronts in tracks or squares configurations propagate much slowly than on controls. (D) Local velocities in the tracks configuration, illustrating that activity propagation along the tracks (vertical direction) is much higher than across them (horizontal direction). Arrows marks tracks with high propagation speed and therefore very small color variation. (E) Box plots of velocity propagation for different fronts of the same culture. On average, propagation along tracks is 18 mm/s, about 6 times larger than across tracks, which is of 3.5 mm/s. **p < 0.01, ***p < 0.001 (Student’s t-test).

We also investigated the velocity of propagating fronts in detail. We considered data at DIV 14 since most fronts encompassed large areas of the 6 mm culture, which provided sufficient statistics for a reliable analysis. The top panels of Figure 4B show representative spatiotemporal fronts for the three configurations, which evolve as quasi-circular fronts from the origin of activity (white circle). As shown in the bottom graph of Figure 4B, their characteristic propagation velocity was obtained by linear regression of *d*(*t*) data (solid lines), where *d* is the distance of each ROI to the origin of activity and *t* its activation time. The measured velocity was *v* ≃ 40 mm/s for controls and substantially decayed to *v* ≃ 13 and 3 mm/s for squares and tracks, respectively. Pearson’s regression coefficients in all three cases were *r* ≳ 0.95. Despite the goodness of the linear regression approach there was, however, a noticeable dispersion in the data for squares and tracks that indicates strong changes in the propagation of the fronts at local scales. For squares, for instance, one could identify a first regime (in the range 0 − 0.2 s) of slow propagation with *v* ≃ 13 mm/s followed by a second one (0.2 − 0.3 s) of fast propagation with *v* ≃ 30 mm/s, which suggests an abrupt change of the underpinned local connectivity during front evolution. For tracks, an inspection of the data revealed that the strong dispersion of some points, with identical time activation of ROIs that were ≃ 6 mm apart, was associated with a much faster propagation of activity along the tracks than transverse to them, as discussed below. The analysis of the velocity using linear regression was consistent across experimental realizations (Figure 4C), with significantly different velocities between controls (⟨*v*⟩ ≃ 42.2 ± 14.7 mm/s), tracks (4.7 ± 1.6 mm/s) and squares (23.4 ± 13.0 mm/s).

The contrast in propagating velocities along PDMS tracks or transversally to them outlined above is analyzed in more detail in Figures 4D-E. For the illustrative spatiotemporal front of Figure 4D we observed that the color scheme along tracks (vertical axis of the image plot) was practically uniform, with characteristic black and red bands (white arrows) that revealed the fast activation of the whole track, typically in a time window on order of 0.3 s. Conversely, the color variation transverse to tracks (horizontal axis) smoothy varied from black at the origin of activity to yellow at the right edge of the culture, leading to a propagation time of 4.3 s. The corresponding velocities were about an order of magnitude dissimilar, with *v*_∥_ ≃ 9.1 mm/s along tracks and *v*_⊥_ ≃ 1.7 mm/s transversally to them. This dissimilarity was preserved across experimental repetitions and was significantly different (Figure 4E, p=0.0027), and on average we obtained ⟨*v*_∥_⟨ = 17.4 ± 9.6 mm/s and ⟨*v*_⊥_⟩ ≃ 3.3 ± 2.2 mm/s.

### Effective connectivity analysis identifies unique network organization traits in topographical cultures

We used transfer entropy to infer causal relations among ROIs in the neuronal cultures (Figure 5), and from them extracted network measures that exposed functional organizational traits. In all cases we used the data presented in the previous figures and at DIV 14. Conceptually, the nodes of the computed effective networks are the ROIs in our experiments, whereas the links are the flow of information among those ROIs. The network traits that we explored include the Global efficiency *G*_*E*_ and the modularity index *Q*. The former captures the capacity of the network to share information as a whole, while the latter informs about the existence of functional modules, *i*.*e*., groups of ROIs that tend to communicate within their group more strongly than with other groups in the network. Figure 5A, top, shows the obtained effective connectivity matrices for the three PDMS configurations, with the modules highlighted as color boxes along the diagonal. In all configurations we observed an abundance of effective connections both within modules and between them. All networks indeed exhibited a similar *G*_*E*_ ≃ 0.45, indicating that neurons in the three configurations easily exchanged information globally. This is understandable in the context of the observed dynamics, in which whole-network correlated activity was present in the three cultures and thus global neuronal communication. The modularity *Q*, however, was clearly different among configurations, with *Q* ≃ 0.26 for controls and *Q* ≃ 0.48 for both tracks and squares. The low *Q* for controls indicates that ROIs within a module connected similarly among themselves and the rest of the network, *i*.*e*., the network effectually operated as a unique system. The high *Q* for the topographical designs indicates the presence of functional microcircuits, *i*.*e*., strongly interconnected neuronal assemblies.

**Figure 5.**
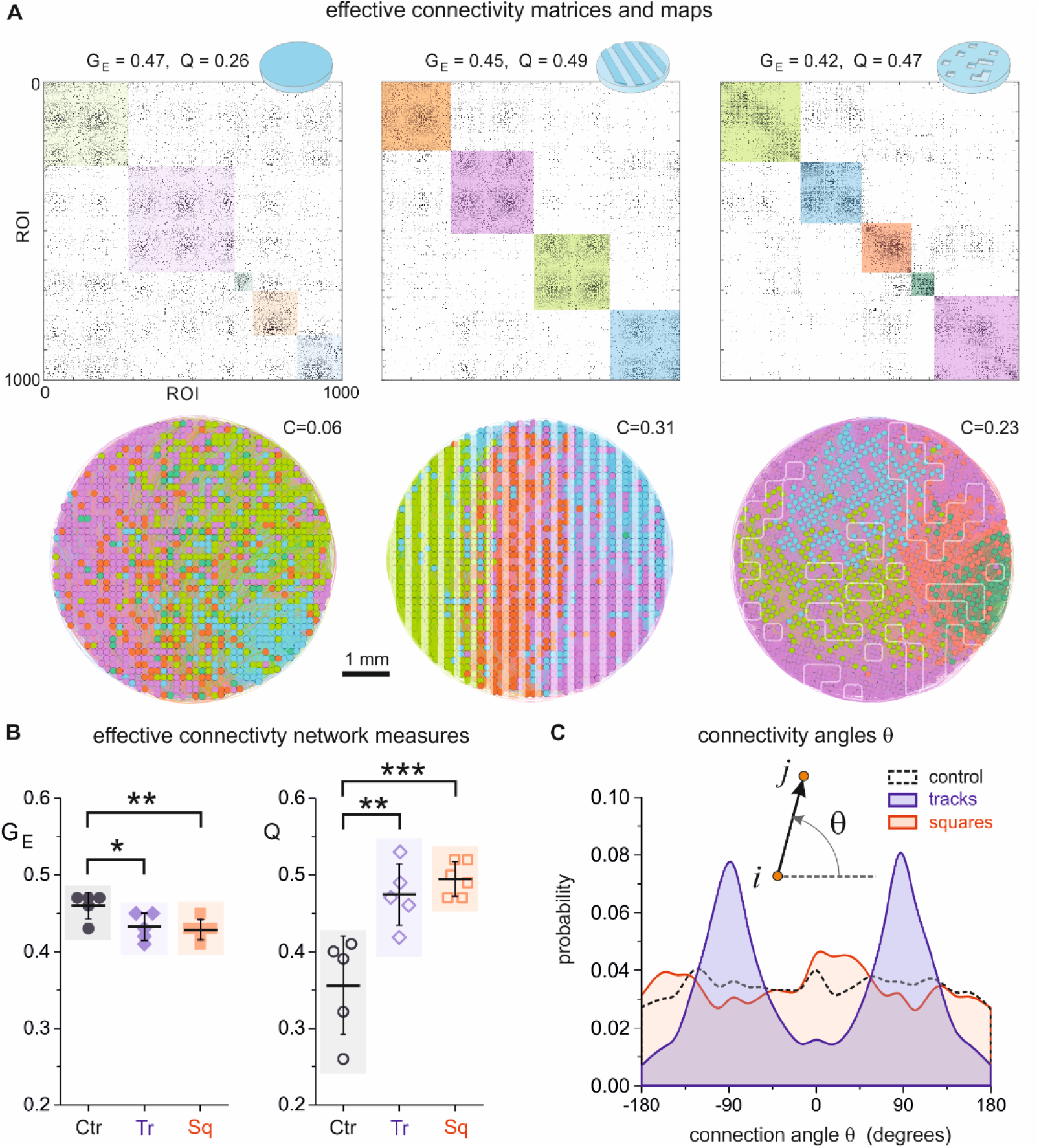
Effective connectivity analysis in PDMS topographical cultures. (A) Top, adjacency matrices of effective connectivity among ROI pairs for the three PDMS configurations. The global efficiency *G*_*E*_ and modularity *Q* values of the networks are indicated on the top. Color boxes along the diagonal of the matrices highlight functional modules, with their color intensity proportional to their strength. Bottom, corresponding network maps, where each dot is an ROI color coded according to the functional module it belongs to. The value of *C* accompanying each map indicates the average spatial compactness of the functional modules, which is markedly high for tracks. (B) Box plots of comparing the network measures *G*_*E*_ and *Q* for the three configurations and including all experimental repetitions. Flat PDMS cultures exhibit a significantly higher *G*_*E*_ and lower *Q* than the other configurations. (C) Distribution of connectivity angles θ, where θ is the angle formed between two effectively connected neurons *ii → jj* and the horizontal axis. The distributions for both controls and squares are approximately flat, while the one for tracks is markedly peaked at 90 and -90 degrees, indicating strong neuronal communication along the tracks. *p < 0.05, **p < 0.01, ***p < 0.001 (Student’s t-test).

The network maps associated to the connectivity matrices (Figure 5A, bottom) provide an additional insight into the impact of PDMS topography on neuronal network communication. These maps show the spatial location of the ROIs in the culture as circles, and color coded according to the functional module they belong to. Effective connections are present, but their abundance masks their individual identification. For controls, the functional modules are spatially interwoven over the area of the culture, a feature that is quantified by the average compactness *C* of the spots and that provided *C* ≃ 0. This mixture of the modules indicates that a neuron tended to interact similarly with any other in the culture and at very long distances. By contrast, for the tracks configuration we observed that functional modules shaped compact spots in the culture (*C* ≃ 0.31) that were aligned with the tracks themselves, indicating that the topographical relief orchestrated the functional organization of the culture. The spatial extent of the functional modules could be related to specific spatiotemporal activity patterns (Figure S3), which indicates a relation between network dynamics and functional traits. A similar compact functional organization was observed for the squares configuration (*C* ≃ 0.23), although the shape of the modules did not concord with the arrangement of the PDMS design possibly due to the high connectivity between top and bottom parts at DIV 14.

The obtained results for functional organization in terms of *G*_*E*_ and *Q* were maintained across experimental realizations (Figure 5B). *G*_*E*_ was not significantly different for the three configurations, but *Q* for tracks and squares was significantly higher than for controls.

To complete the analysis, we investigated in more detail the emergence of local functional features in the topographical cultures. Given the capacity of tracks to funnel neuronal connectivity along their length, we examined whether such a privileged direction was captured by the effective connectivity analysis. For that, we computed the angle of effective connections relative to the horizontal axis and plotted the distribution of angles. As shown in Figure 5C, controls (black) showed a homogenous distribution of angles that indicated an isotropic effective connectivity. A similar result was observed for the squares configuration (red), although the distribution exhibited stronger fluctuations. For tracks (blue), however, neat peaks appeared at -90 and 90 degrees, distinctly revealing a preferred vertical direction in neuronal communication. Thus, the effective connectivity analysis shown here demonstrated to be an invaluable tool to complete the activity and immunostaining analyses, and brought to light additional evidence that the PDMS relief help dictating the way in which neurons wired and communicated.

## DISCUSSION

Primary neuronal cultures are one of the most celebrated techniques in several multidisciplinary research fields, including physics of complex systems, neuroengineering and medicine. Their versatility, accessibility and ease of manipulation have made them ideal to investigate in a controlled manner phenomena as diverse as self-organization, repertoire of activity patterns, structure-to-function relationship, resilience to perturbations and alteration upon disease. However, primary cultures grown on flat surfaces typically exhibit a strong bursting behavior in which all neurons activate together in a short time window and remain practically silent in between bursts. In the present study we showed that this all-or-none rigid behavior can be relaxed by incorporating spatial anisotropies on the substrate, in the form of topographical reliefs, that substantially enrich the repertoire of activity patterns and approach them to what is observed *in vivo*, where activations of different spatiotemporal structure coexist.

The spatial constraints promoted by the PDMS reliefs favored local connectivity and facilitated activity at a microcircuit level, but without suppressing whole-network dynamics. Thus, our design promoted the emergence of neuronal networks with balanced integration and segregation, *i*.*e*., where both local computation and whole-network communication coexisted. Other studies investigated the realization of this balance by using modular designs (Park et al., 2021; Shein-Idelson et al., 2011; Yamamoto et al., 2018), in which neurons were spatially confined in small areas. Our PDMS topographical modulation provides an alternative approach to such an *ad hoc* confinement, shaping dynamically rich networks in both space and time without fine tuning the position of neurons and axons. We note that a broader modulation of the dynamical repertoire in our cultures could be achieved by altering the ratio between excitatory of inhibitory neurons or by decreasing the strength of excitatory synapses. All our experiments were conducted with both excitation and inhibition active. Given the role of inhibition in modulating activity (Isaacson and Scanziani, 2011; Sukenik et al., 2021), we argue that the repertoire of activity patterns could additionally be tuned by blocking GABA receptors in inhibitory neurons, which would make bursting events more similar among themselves, or by reducing the amount of excitatory transmission by blocking AMPA-glutamate receptors. The latter was explored for instance in modular networks (Yamamoto et al., 2018), observing that the integration-segregation balance shifted towards higher segregation as excitation was reduced.

We found a PDMS height of around 70 µm to be optimal, since it promoted a coarse axonal positioning and orientation in a neuronal neighborhood, *e*.*g*., along a PDMS track, while allowing for easy interconnection with other neighborhoods. Our observation that connections follow the track ridges is in agreement with other studies that investigated in detail the impact of geometrical cues on axonal growth (Basso et al., 2019). PDMS heights of 100 µm or higher in our experiments often shaped isolated regions in the culture, whereas heights of ∼30 µm did not cause sufficient structural alterations to markedly modify network dynamics. Other studies also used PDMS reliefs to control connectivity in neuronal circuits, most notably holes and pillars of characteristic scales in the range 10 − 100 µm (Li et al., 2017, 2014), ultrasoft PDMS (Sumi et al., 2020), or by combining fine-tuned neuroengineering and microfluidics (Forró et al., 2018; Holloway et al., 2021; Liu et al., 2021; Marconi et al., 2012; Nikolakopoulou et al., 2021). In these works, authors reported a richer repertoire of activity patterns or the suppression of extreme bursting. In the context of these studies, the relevance of our PDMS reliefs is that cells’ neurite development is coarsely guided instead of fully delineated, allowing the circuit to retain its self-organization potential. The effort of imprinting ‘mesoscale architecture’ while allowing self-organization is conceptually similar to the studies on neuronal cultures with spatial aggregation (Okujeni et al., 2017; Okujeni and Egert, 2019; Tibau et al., 2020). Aggregation helped neurons to connect within their neighborhood but without hindering long-range connectivity, shaping networks with a richer dynamic behavior and more varied activity initiation, as in our case.

The contrasting velocities of activity propagation in the tracks configuration, with measured velocities about 5 times larger along tracks than across them, can be put in context of theoretical models (Bressloff, 2000; Golomb and Ermentrout, 1999) and experiments on activity propagation in one-dimensional neuronal cultures (Feinerman et al., 2005; Jacobi et al., 2010; Jacobi and Moses, 2007). These studies revealed that the velocity of activity fronts depended on average neuronal connectivity and synaptic strength. If we assume that synaptic strength is similar in all neurons in our cultures, then we conclude that connectivity along tracks was 5-fold higher than transverse to it. This difference is consistent with the functional data of Fig. 5C, in which effective connections along tracks was about 8 times more abundant than across tracks. However, it must be note that effective connectivity reflects communication and not synaptic links, and therefore a direct quantitative analysis of structural connectivity is not possible based solely on the analysis of the functional data. We also remark that the maximum velocity that we measured along tracks was about 30 mm/s, which is slower compared to the data provided in the study of Feinerman and coworkers (Feinerman et al., 2005), who measured velocities in the range 40-80 mm/s. In Feinerman’s study, however, activity was solely driven by excitatory neurons, while in our networks both excitation and inhibition are active. Thus, the presence of inhibition is possibly the reason for the comparatively low velocities measured in our experiments.

Given the importance of spatial constraints in molding neuronal circuitry architecture and function (Stiso and Bassett, 2018), it is interesting as a future exploration to investigate the impact of different degrees of connectivity restrictions. Chemical patterning (Liu et al., 2021; Yamamoto et al., 2018) offers the delineation of precise circuits but with restricted self-organization capacity, while PDMS topographical modulation offers a broader flexibility at an expense of a poorer control on physical wiring. We observed in our experiments that different repetitions of the tracks or squares designs led to neuronal circuits with similar global dynamic behavior but with different functional details. For instance, the distribution of initiation points (Fig. 4) and functional modules (Fig. 5) varied across repetitions, indicating different mesoscopic evolution. Additionally, some of the explored cultures tended to become more integrated and with increasingly stronger network-wide bursting as they matured, suggesting that the initially imprinted inhomogeneities were erased at long term. This loss of richness was particularly strong in the squares configuration. We hypothesize that, to better approach brain-like behavior *in vitro*, an optimal experimental system would be one that combines topographical and chemical patterning, thus preserving key functional traits without the loss of flexible self-organization. We also conjecture that external stimulation, e.g., as in (Poli et al., 2016), may be a necessary ingredient to shape circuits with long-lasting functional features.

Finally, we note that our experiments may be of interest for those studies that use neuronal cultures as models for neurological disorders *in vitro*. These studies often explore the alterations in network collective activity caused by a disease. For instance, in a recent study of Parkinson *in vitro* (Carola et al., 2021), authors observed that the affected networks exhibited a much higher number of network-wide bursts as compared to healthy controls. Although their results were conclusive, the investigation was difficulted by the tendency of standard, glass-grown neuronal cultures to exhibit persistent whole-network bursting. Thus, we argue that the use of PDMS topographical substrates may help to prepare networks whose activity is much varied since early development, identify the impact of a disease in network formation, activity and functionality, as well as their evolution along time.

### Limitations of the study

Neurons in our experiments were plated in a homogeneous manner in PLL-coated PDMS surfaces. However, we often observed that neurons strongly aggregated after few days, or that some areas of the PDMS relief, generally at the bottom, were not occupied by cells. This problem was present both in the tracks and squares configurations, and often led to large empty areas for the latter. We ascribe this lack of homogeneity to the PLL coating, which possibly was not sufficiently uniform for neurons to adhere to the surface, or to capillary forces that caused the trapping of air bubbles and blocked coating. These inhomogeneities can be observed for instance in the fluorescence image of the tracks configuration in Figure 1. We note that fluctuations in local density accentuated the anisotropies induced by the relief and, in turn, amplified the variability of the spatiotemporal fronts. Specifically, for the tracks configuration, the distinct parallel and transverse velocities is possibly favored by the contrasting neuronal densities between top and bottom parts. Nonetheless, we used high-resolution phase contrast images and immunostaining to reject cultures in which neurons grew as isolated patches. All studied cultures here contained neurons that were globally interconnected and exhibited episodes of coordinated activity that encompassed from few neurons to the entire network.

On the other hand, in this study we were interested in the collective behavior of mm-sized cultures rather than in the precise individual dynamics of their constituting neurons. The need to access a large field of view in combination with limitations in image resolution imposed by the fluorescence camera, made not possible to resolve single cells. Hence, we analyzed network activity using an ROI approach. To investigate whether this approach could create artifacts, we run experiments in smaller, 4 mm diameter cultures in which both single neuron monitoring and ROIs could be used (Orlandi et al., 2013; Tibau et al., 2020). Similar qualitatively results were obtained when comparing both approaches for all major dynamic and functional descriptors. Thus, the ROIs analysis can be viewed as a coarse-graining approach, which suffices to capture interesting mesoscale phenomena as far as the spatial extent of these phenomena is larger than the characteristic neuron size. In our case, network bursts and functional modules covered areas on the order of few mm, much larger than the 10 *µm* diameter of a neuron. We believe that this coarse-graining may be a source of inspiration to explore neuronal circuits at different scales and may help bridging the gap between *in vitro* networks and naturally formed neuronal circuits. Related to this, it is important to emphasize that we considered rat primary cortical cultures for our experimental design. Alternative cells models such as human induced pluripotent stem cells (hiPSCs) may broaden the spectrum of dynamical and functional traits shown here. Indeed, hiPSCs cultures grown on flat substrates already exhibit a higher individual activity and a richer repertoire of coordinated activations (Carola et al., 2021; Kirwan et al., 2015), as opposed to rat primary cultures that show a strongly rigid bursting behavior.

## METHODS

All methods can be found in the Transparent Methods section of the Supplemental Information file.

## SUPPLEMENTAL INFORMATION

Supplemental Information includes a detailed description of the methods, videos of representative recordings, and additional analyses of experimental data.

## ACKNOWLEDGMENTS

This work was funded by European Union’s Horizon 2020 project MESO-BRAIN, Grant 713140; by the Ministerio de Ciencia e Innovación (Spain), Grant PID2019-108842GB-C21; and by the Generalitat de Catalunya, Grant 2017-SGR-1061. The authors would like to acknowledge Hideaki Yamamoto (Tohoku University, Sendai, Japan) and Sergio Faci-Lázaro (University of Zaragoza, Spain) for highly valued scientific discussions.

## AUTHOR CONTRIBUTIONS

J.S. and M.M.-F. designed the experiments. M.M.-F. fabricated the topographical substrates and prepared the neuronal cultures. M.M.-F. and C.F.L.-L. performed the experiments. J.S. and M.M.-F. developed the software for data analysis. J.S., M.M.-F., C.F.-L. and T.F. designed and performed the analyses. D.T. carried out the immunohistochemical analyses. T.F., P.M. and S.M. contributed to analysis tools. J.S. and M.M.-F. prepared the original figures. J.S. and M.M.-F. wrote the manuscript. J.S. acquired funding to conduct the research. All authors have read, corrected, and approved the final version of the manuscript.

## DECLARATION OF INTERESTS

The authors report no competing interests.

## Transparent Methods

### PDMS topographical reliefs

Topographical substrates were prepared by using a specially designed printed circuit board (2CI Circuitos Impresos, Spain) that served as a negative mold for the desired topographical design. As shown in Figure S1A, the printed circuit was formed by two layers, a bottom one of uniform fiberglass 2 mm thick and a top one of cooper deposits 70 µm high that shaped different designs. The height of the copper was constant along the board. For the present work, two main designs were used and termed ‘tracks’ and ‘squares’ (Figure S1B). ‘Tracks’ consisted of parallel rectangular bands 300 µm wide and 20 mm long and separated by 300 µm. ‘Squares’ consisted of randomly positioned square blocks of 300 µm lateral size. Blocks were placed following a grid of 300 µm spacing, so that there was no overlap between blocks and the spatial dimensions of the resulting designs were all multiple of the basic square dimensions. The blocks were laid on a 20×20 mm^2^ area and occupied 15% of it. PDMS (Sylgard 184, Farnell) with a mixture of 90% base and 10% curing agent was poured on the printed circuit board and cured at 90 °C for 2h. The PDMS was then gently removed, shaping a topographical relief in which the copper and fiberglass on the board corresponded to depressions and crevices on the PDMS, respectively (Figure S1C).

PDMS discs 6 mm in diameter and typically 1 mm thick were pierced using stainless steel punchers (Bahco 400.003.020) and then carefully washed, dried, and attached to glass coverslips (#1 Marienfeld-Superior). A coverslip accommodated two PDMS discs (Figure S1D). Different sets of coverslips containing PDMS were prepared and sterilized in an autoclave (Selecta 4002515), which in turn strongly bond the PDMS to the glass surface. A detail of the ‘Tracks’ and ‘Squares’ designs ready for culturing are provided in Figure S1E. The bright-field images are accompanied of simple sketches to clarify which are the top and down areas of the designs. Since the copper in the printed board is very smooth, the corresponding PDMS depressions were transparent when observed under bright-field microscopy, whereas the slight roughness of the fiberglass led to PDMS crevices that appeared opaquer. These slight differences in opacity did not affect calcium fluorescence imaging on neurons over the relief. To compare neuronal network dynamics with and without topography, flat PDMS discs were also prepared on plastic Petri dishes and pierced with the aforementioned punchers. These neuronal cultures are referred as ‘controls’ in the main text.

### Primary neuronal cell cultures

The preparation of primary neuronal cultures was carried out in accordance with the regulations of the Ethical Committee for Animal Experimentation of the University of Barcelona (approved ethical order DMAH-5461) and the laws for animal experimentation of the Generalitat de Catalunya (Catalonia, Spain).

In all experiments, primary neurons from Sprague-Dawley embryonic cortices at days 18-19 of development were used and following identical protocols as described previously (Orlandi et al., 2013; Tibau et al., 2020, 2013). Rats were provided by the animal farm of the University of Barcelona. Cortices’ dissection and cell culturing were conducted at the laboratory of Dr. Soriano at the Faculty of Physics of the University of Barcelona. Briefly, dissection was carried out in ice-cold L-15 medium (Gibco), cortical tissue dissociated mechanically by repeated pipetting, and neurons suspended in plating medium [90% Eagle’s Minimum Essential Medium (MEM, Invitrogen) with 5% Horse Serum (HS, Invitrogen), 5% Bovine Calf Serum (Invitrogen) and 1 µl/ml B27 (Sigma)]. Prior culturing, PDMS surfaces were submerged overnight in the adhesive protein Poly–L–Lysine (PLL, Sigma-Aldrich) at a concentration of 10 mg/ml PLL in Borate Buffer. Neurons were plated on these PDMS surfaces, procuring cultures with a neuronal density of about 400 neurons/mm^2^. Although neurons were homogenously distributed over the surface, they tended to slightly aggregate. One day after plating, cultures were infected with adeno-associated viruses bearing the GCamp6s calcium sensor under synapsin-I promoter (AAV9.Syn.GCaMP6s.WPRE.SV40, Addgene). Four days afterwards, at day *in vitro* (DIV) 5, plating medium was replaced by changing medium (90% MEM, 10% HS and 0.5% FUDR) to limit glial growth. At DIV 8 the medium was switch again to final medium (90% MEM and 10% HS) and refreshed periodically every 3 days.

Only neurons express the calcium sensor under the CaMKII promoter. Thus, although cultures contained both neurons and glia, only neuronal activity was visualized.

4 wells (8 PDMS cultures) were prepared in each dissection and were kept in 4-well plates (Nunc), each plate hosting 2 wells. This facilitated the consecutive recording of different cultures while minimizing possible alterations in those wells that were not recorded at that moment. Cultures were incubated at 37 °C, 5% CO_2_, and 95% humidity. Spontaneous activity emerged by DIV 5, but GCAMP6s expression was not sufficiently strong for reliable imaging until DIV 7.

All cultures contained both excitatory and inhibitory neurons, with approximately 80% excitation and 20% inhibition (Soriano et al., 2008).

### Immunocytochemistry

Neuronal cultures were fixed with 4% PFA (Sigma) for 15 min at room temperature, rinsed with PBS and incubated with blocking solution containing 0.03% Triton (Sigma) and 5% Normal Donkey Serum (Jackson Immunoresearch) in PBS for 45 minutes at room temperature. Primary antibodies against neuronal cytoskeleton (β3-Tubulin and MAP2, Table 1) were applied diluted in blocking solution and incubated overnight at 4°C. Alexa488-conjugated secondary antibody against mouse (Sigma, A10667) was diluted in blocking solution and incubated for 90 minutes at room temperature. For astrocytic staining, samples were post-fixed again and incubated with anti-GFAP antibody directly conjugated with Cy3 (Table 1) overnight at 4°C. Then, cultures were rinsed with PBS and mounted using DAPI-fluoromount–G (ShouternBiotech). Immunocytochemical images were acquired on a Zeiss confocal microscope (LSM-880).

**Table 1:** Primary antibodies

| Antigen | Company | Origin | Code |
| --- | --- | --- | --- |
| Tuj1 | Sigma | Mouse | T8660 |
| MAP2 | Sigma | Mouse | M1406 |
| GFAP-Cy3 | Sigma | Mouse | C9205 |

### Calcium imaging

Spontaneous activity in neuronal cultures grown on the PDMS topographical substrates was recorded daily at 25°C along two weeks, from DIV 7 (the onset of strong GCAMP6s calcium signal) to 21 (the beginning of culture degradation). Since monitoring the same culture was crucial, the set of prepared wells was inspected in detail before any recording, and cultures that did not have a homogenous distribution of neurons or that were inactive at DIV 7 were discarded.

For the selected cultures, the 4-well plate in which they sit was mounted on a Zeiss Axiovert C25 inverted microscope equipped with a high-speed CMOS camera (Hamamatsu Orca Flash 4.1). The combination of a 2.5X objective and an optical zoom allowed for the visualization of an entire 6 mm culture with a spatial resolution of 5.9 µm/pixel, an image size of 1,024×1,024 pixels, and 8-bit grey scale format. Spontaneous activity recordings were carried out for 30 min at 100 images/s and repeated every 24h. The orientation of a given culture relative to the camera was maintained along the 2-week culture evolution to facilitate data analysis.

In total, the number of repetitions (complete series of daily monitoring on the same culture) were 5, 5, and 6 for control, tracks, and squares, respectively.

### Calcium imaging data analysis and activity events detection

Fluorescence recordings were analyzed with the custom-made software Netcal run in Matlab (Fernández-García et al., 2020; Orlandi et al., 2017). For convenience, and since the scope of the study was to investigate collective behavior, a set of 1,300 Regions of Interest (ROIs) were laid out on the image (Figure S2A). The ROIs shaped a grid centered at the culture and extending its entire circular shape of 6 mm diameter. A ROI had a typical size of 150×150 µm and contained about 5-10 neurons. The average fluorescence intensity of each ROI *i* along the 30 min recording duration was then extracted, and the obtained fluorescence trace *F*_*i*_ (*t*) was corrected from drifts and normalized as Δ*FF*_*i*_ ≡ (*F*_*i*(*t*) −_ *F*_*i*,0_)/*F*_*i*,0_, where *F*_*i*,0_ is the basal fluorescence level (without neuronal activity) (Figure S2B).

Fluorescence data of each ROI was converted into time series of neuronal activity by using the Schmitt trigger method, which accepts a sharp change in fluorescence as an activity episode whenever the fluorescence stays elevated for at least 100 ms between a lower and a higher threshold (Grewe et al., 2010). The two thresholds were necessary to prevent camera noise or other artifacts to be identified as activity events. The Schmitt trigger method captured the onset time of activity in each ROI, independently on the amplitude of the fluorescence peak as far as it was sufficiently high. The train of detected events, extended to all ROIs in the network, was visualized as raster plots (Figure S2C) and framed the core dataset for in-depth analysis of the neuronal cultures.

### Population activity, network bursts, and distribution of burst amplitudes

The population activity *A* quantified the capacity of the neurons in the network to exhibit coordinated activity. It was computed as the fraction of ROIs in the network that activated together without repetition in a sliding window 1 s wide and 0.1 s step. *A* varied between 0 (no activity) and 1 (full network activation). Sharp peaks in A identified strong coordinated activity and were denoted as *network bursts*. These bursts were deemed significant when their amplitude *A*_*b*_ verified *A*_*b*_ > μ_*bgnd*_ + 3 · *SD*_*bgnd*_, where μ_*bgnd*_ and *SD*_*bgnd*_ are the mean and standard deviation of background activity. In general, for most of the experiments, significant bursts were those with *A*_*b*_ > 0.1, *i*.*e*., 10% of the network active. All significant burst amplitudes *A*_*b*_, across repetitions and for a given experimental condition, were pooled together to build the distribution of amplitudes *A*_*b*_. These distributions were finally compared between the different topographical designs and the control, flat PDMS condition.

### Dynamical richness

The dynamical richness Θ provides a measure of the spatiotemporal variability of network activity, *i*.*e*., the existence of a broad range of coactivation patterns and dynamical states, and is defined as (Yamamoto et al., 2018):

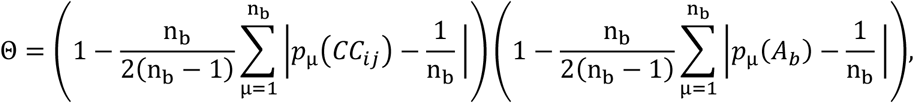

where *p*(*C*_*ij*_) is the distribution of Pearson’s correlation coefficients between the activity trains of all pairs of ROIs *i* and *j, p*(*A*_*b*_) the distribution of burst sizes *A*_*b*_, | · | denotes the absolute value and n_b_ = 20 is the number of bins used for estimating the distributions. Θ varies between 0 (no richness) and 1 (full richness). Conceptually, Θ ≃ 0 corresponds to a scenario of random activity or coherent, whole-network activations, whereas Θ ≃ 1 corresponds to a network state in which neurons coactivate in groups of richly varying size and temporal occurrence.

### Bursts’ analysis as spatiotemporal fronts

Network bursts propagated throughout the PDMS surface as spatiotemporal fronts whose structure and velocity depended on the underpinned topographical design. The propagation of a given burst was depicted in Figure 2A as an image plot, in which each active ROI was shown in the Euclidean *x* − *y* space and colored according to its activation time. Dark colors represented those regions that activated first, and yellow-white colors those that activated the latest. Inactive ROIs or regions of the map without ROIs were shown in dark grey. The origin of activity, termed ‘burst initiation point’, was computed by considering a group of 10 ROIs with the shortest activation times and by analyzing all combinations of 4 ROIs within the group, computing for each combination the average inter-ROI Euclidean distance *d*_0_ and average activation time *t*_0_. The combination that procured the lowest *d*_*0*_ and shortest *t*_0_ (termed 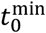) was selected as initiator and the ROIs’ centroid (*x*_0_, *y*_0_) was evaluated. This centroid was ascribed as the burst initiation point and was shown in the image plots of Figure 2A as a white circle. Finally, the activation times *t*_*i*_ of all ROIs *i* were then shifted according to the origin of activity as 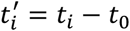. ROIs with negative time 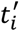 values were set to 0.

The collection of burst initiation points for each experiment was further analyzed to study their spatial distribution and quantify the tendency for the spontaneous activity to start in the same area of the culture. Following (Faci-Lázaro et al., 2019; Orlandi et al., 2013), the distribution of points was converted to a probability density function of burst initiation, from which the Gini coefficient *λ* was extracted as a measure of activity focalization. *λ →* 1 indicated a tendency towards a strongly focalized initiation in the same spot, whereas *λ →* 0 indicated a tendency towards a homogeneous distribution of initiation across the culture.

### Similarity of spatiotemporal fronts

The richness of activity repertoire in a culture was quantified by analyzing the similarity among bursts’ spatiotemporal structure. A coarse approach (Figure 2B) consisted in plotting the bursts as blue-red color bands, where the width of the band is the number of active ROIs and its color patterning indicates the propagation of activity. Each band has its own ROI ordering, from the one that started activity to the last. Bursts with similar band widths and color schemes indicate a comparable propagation structure, *i*.*e*., similar number of active ROIs and temporal evolution. A more refined approach was carried out as follows. First, each burst *i* was treated as a vector *v*_*i*_ whose elements contained the activation time of each ROI. Non-active ROIs were set to −1. All bursts’ vectors preserved the same indexing of ROIs, *i*.*e*., a given position in all vectors contained the activation time of the same ROI. Second, for each pair of bursts’ vectors *v*_*i*_ and *v*_*j*_, the number of common active ROIs (non-negative entries in both vectors) was determined, and this number was divided by the burst that had the largest number of active ROIs. This procured a ‘common ROIs’ matrix ***C*** = *c*_*ij*_, in which bursts that shared most of the ROI indexes had *c*_*ij*_ *→* 1, otherwise *c*_*ij*_ *→* 0. Third, cross correlation was carried out between all pairs of bursts’ vectors but using only those ROIs that were active in both vectors, leading to a correlation coefficients matrix ***R*** = ***r***_*ij*_. Values of ***r***_*ij*_ *→* 1 indicated pairs of bursts with almost identical spatiotemporal structure (same active ROIs and propagation times), while r_*ij*_ → 0 indicated bursts that shared few ROIs or that propagated in a completely different way. A final matrix **S** of similarity among burst pairs was obtained as the element-wise multiplication of ***C*** and ***R***, *i*.*e*., ***S*** = ***C* ∘ *R***. To classify the bursts and visualize the matrix ***S***, community structure analysis was carried out in ***S*** using a fast implementation of the Louvain algorithm (Blondel et al., 2008), procuring a new matrix whose elements were ordered according to the detected communities (Figure 2C). For clarity of data visualization, bursts’ indexes in Figure 2B were also sorted according to the detected communities in Figure 2C.

### Velocity of propagating fronts

The propagation speed of activity fronts was analyzed by computing the Euclidean distance ρ_i_ of each ROI *i* to the origin of activity (*x*_0_, *y*_0_) and by plotting next ρ_i_ as a function of 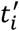, the activation times relative to the origin of activity. An estimation of the global propagation velocity was obtained as the slope of a linear fit 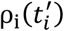, and with the intercept forced at (0,0). Only fits with Pearson’s regression coefficients ***r*** ≥ 0.95 were accepted, with the best fits corresponding to bursts that propagated as neat circular fronts (Orlandi et al., 2013) and that were typically observed in control cultures. The determination of parallel and transverse velocities for the ‘tracks’ configuration was carried out similarly but after selecting either a row of ROIs in the culture (parallel velocity) or a column (transverse velocity).

### Effective connectivity

Causal relationships among pairs of ROIs’ activity were computed by using a modified version of Generalized Transfer Entropy (GTE) (Ludl and Soriano, 2020; Stetter et al., 2012; Tibau et al., 2020) run in Matlab. Binarized vectors for the 30 min activity trains (‘1’ for the presence of a spike, ‘0’ for absence) were constructed using a time bin of 20 ms, and an effective connection from ROI to ROI *J* (TE_*I→*J_) was established whenever the information contained in *I* significantly increased the capacity to predict future states of *J*. Instant feedback was present, and Markov Order was set to 2 (Stetter et al., 2012). The significance threshold *z* for effective connections was established by comparing the transfer entropy estimate TE_*I→*J_ with the joint distribution of all input *X* to *J* and output *I* to *Y* (for any *X* and *Y*), as

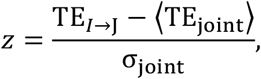

where ⟨TE_joint_⟩ is the average value of the joint distribution and σ_joint_ its standard deviation. Significant connections were then set as those with z-score *z* ≥ 2. This threshold was considered optimal to capture effective communication both at global and local scales (Ludl and Soriano, 2020), *i*.*e*., whole-network collective activity and interactions at the PDMS relief level. Significant TE scores were finally set to 0 (absence of connection) or 1 (connection present), shaping directed yet unweighted connectivity matrices. These matrices were visualized in the form of network maps with Gephi (Bastian et al., 2009).

### Network measures

Effective connectivity matrices were analyzed using the ‘Brain Connectivity Toolbox’ (Rubinov and Sporns, 2010), run in Matlab, to quantify their topological organization. Two network measures were used here, namely the *global efficiency G*_*E*_ (Latora and Marchiori, 2001) and the *modularity index Q* (Blondel et al., 2008).

*G*_*E*_ accounts for the capacity of neurons to exchange information across the entire network, and is defined as

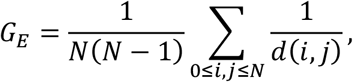

where *N* is the number of ROIs and *d*(*i, j*) is the length of the shortest topological path connecting ROIs *i* and *j*, with non-connected ROIs procuring *d*(*i, j*) = ∞. *G*_*E*_ ≃ 0 indicates that any ROI poorly communicates with any other in the network, while *G*_*E*_ ≃ 1 indicates that there is a strong capacity for information exchange at the whole-network scale.

The modularity index *Q* accounts for the tendency of neurons to form functional modules, *i*.*e*., groups of neurons that are more connected within their groups than with neurons in other groups, and is defined as

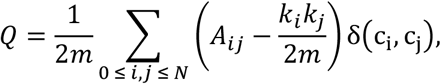

where *N* is the number of ROIs, *A*_*ij*_represents the weight of the connection between *i* and *j*, 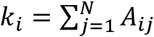 is the sum of the weights of the connections attached to neuron *i, c*_*i*_ is the community to which neuron *i* belongs, 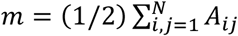, and *δ*(*u, v*) is the Kronecker Delta with *δ*(*u, v*) = 1 for *u* = *v* and 0 otherwise. Optimal community structure was computed using the Louvain algorithm (Blondel et al., 2008). *Q* ranges between 0 (the entire network shapes a unique module) and 1 (each ROI is an isolated module). Values of *Q* ≳ 0.3 indicate the clear existence of modules in the network and interconnected to one another.

The robustness of the community structure analysis was double-checked using the ‘stochastic block model’ (Peixoto, 2017), which infers community structure using a combination of a generative process and a reduction process that minimizes the amount of information required to describe the network. The generative process is based on the notion of groups of nodes: it partitions the nodes into groups and counts the edges in and across groups, such that nodes that belong to the same group possess the same probability of being connected with nodes of another group in the network. The precise implementation used is that of the graph-tool library (Peixoto 2014). The overall structure of the inferred modules was very similar to those obtained with the Louvain algorithm, although the precise number and size of the modules slightly varied. As in Louvain’s approach, the ‘block model’ rendered modules that were spatially compact, followed the substrate topography, and captured by themselves the dominant spatiotemporal fronts.

### Spatial compactness of functional modules

Compactness *C* refers to the property of objects to exhibit a minimum perimeter *P* for a given area *S*, and is mathematically measured through the Polsby-Popper test, *C* = 4π*S*/*P*^2^, with *C* = 0 for a lack of compactness, *e*.*g*., randomly scattered spots, and 1 for a circle, the most compact shape. The compactness of the effective networks shown in Figure 5 was determined as follows. For each functional module, its participating ROIs were drawn as solid white squares on a black background. ROIs were laid down following a grid, so that two adjacent ROIs shaped a solid rectangle. The final white object containing all participating ROIs was then processed to eliminate single black squares surrounded by white regions. This was necessary to prevent that few empty regions could dominate the perimeter of the object. The compactness *C*_*i*_ for the object (functional module) *i* was then computed. To correct for the artifact associated with the square shape of the ROIs, which increased the perimeter of the object and procured lower compactness than expected, a reference compactness for the entire culture *C*_culture_ was also determined by using all original ROIs of the experiment and that, by construction, shaped a circle. Typically, *C*_culture_ ≃ 0.65, smaller than the expected value of 1 associated to a perfect circle. Thus, for each functional module, its compactness was corrected as 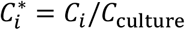. The compactness values shown in Figure 5 were finally obtained, for a given culture, as 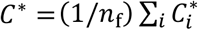, with *n*_*f*_ the number of functional modules.

## Supplementary Videos

**SV1 (DIV14 Tracks 2 min):**

Spontaneous activity for the tracks configuration along 2 min. The video plays at approximately 5x real acquisition time for a better appreciation of the activity fronts and their propagation. The video is related to Figs. 1 and 2.

**SV2 (DIV14 Tracks 20 min):**

Same video as before but extended to 20 min to observe the rich repertoire of activity patterns. Playback is 40x acquisition time. Related to Figs. 1 and 2.

**SV3 (DIV07 Squares 2 min):**

Spontaneous activity for the squares configuration along 2 min. The video plays at approximately 5x real acquisition time. Related to Figs. 1 and 2.

**SV4 (DIV14 Tracks detail 2 min):**

Spontaneous activity in a zoom-in region of the tracks configuration. Playback is 30x. At approximately recording time 54 s the processes (neurons and dendrites) in the culture can be appreciated. Several of these processes are aligned along the left track.

**SV5 (DIV14 Squares detail 2 min):**

Corresponding zoom-in video for the squares configuration. Playback is 9x.

## Supplementary Figures

**Figure S1.**
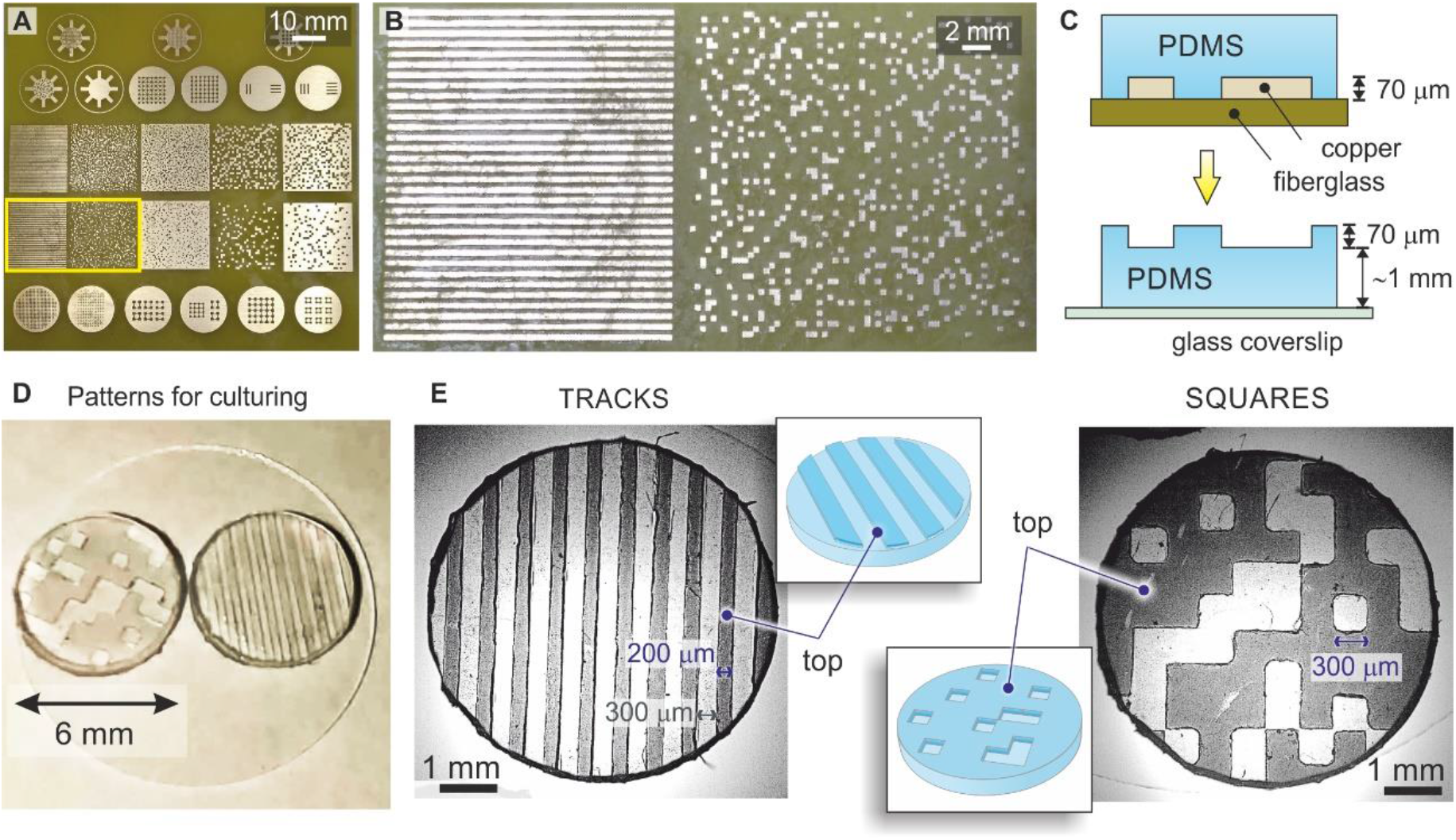
Preparation of topographical PDMS substrates. (Related to Figure 1)

(A) Printed circuit board containing different designs. Copper (bright goldish color) elevates over the fiberglass (dark green) by 70 µm. The yellow rectangle highlights the designs used in the present study. (B) Detail of ‘tracks’ (left) and ‘squares’ (right) designs. For tracks, copper tracks elevations are 200 µm thick and are separated by crevices 200 µm wide. For squares, they are 300×300 µm unit size and randomly positioned without overlap and occupying 15% of the available area. (C) Sketch of PDMS casting from the printed circuit board. (D) A couple of topographical cultures 6 mm in diameter attached to a glass coverslip. (E) Detailed bright-field images of the topographical patterns ready for cell culturing. Transparent and dark areas correspond to PDMS in contact during curing with copper or fiberglass, respectively, and shape the bottom and top regions of the topographical relief.

**Figure S2.**
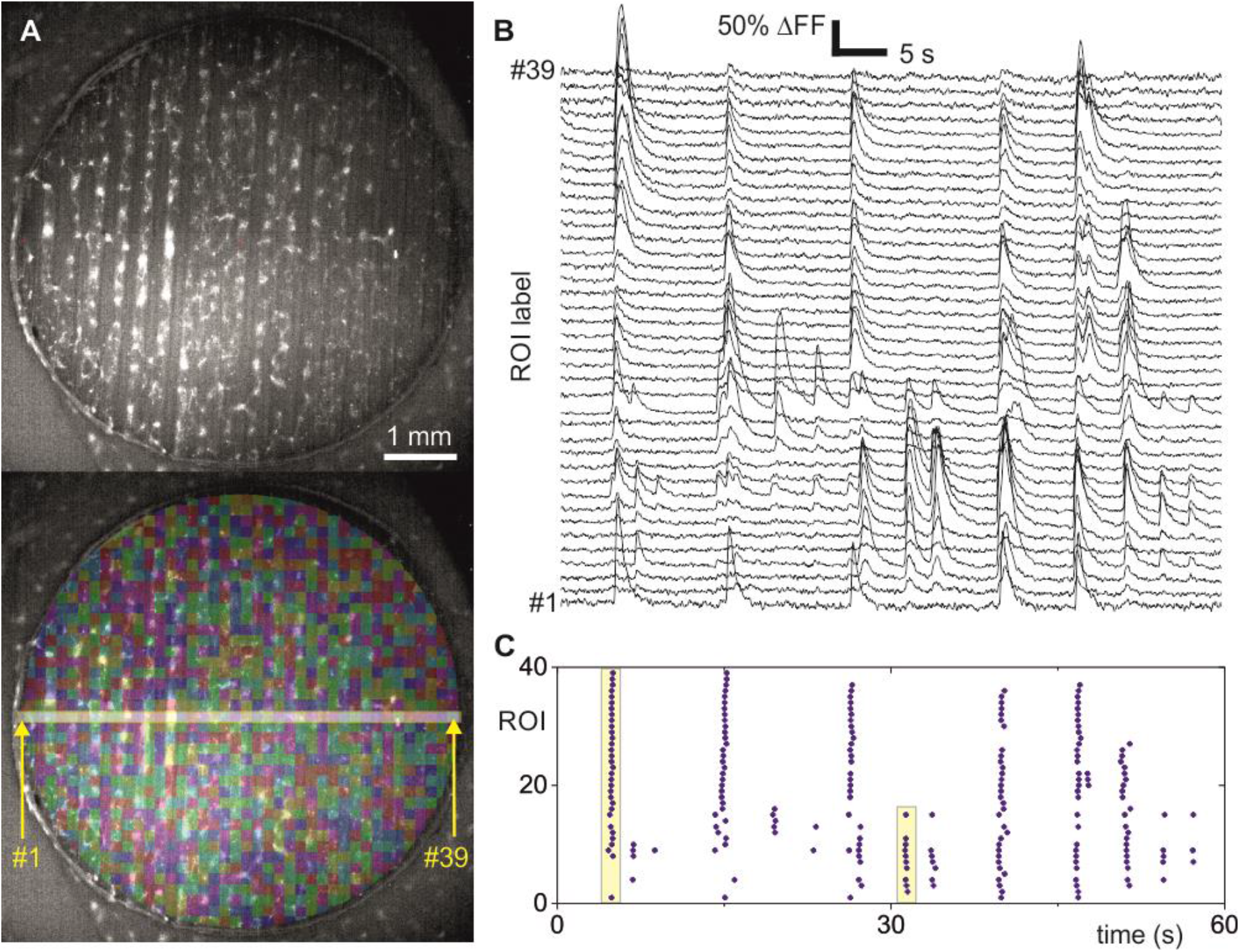
Data analysis. (Related to Figure 1)

(A) Top, highly contrasted fluorescence image of a ‘tracks’ culture. Bottom, Regions of Interest (ROIs) set as squared boxes and covering the entire circular culture, with a total number of 1,300. The 39 ROIs at the center of the culture are used to provide representative fluorescence traces. (B) Normalized fluorescence traces for the 39 central ROIs along the first minute of the recording, vertically shifted for clarity. Sharp peaks indicate neuronal activity. (C) Corresponding raster plot. Examples of coordinated neuronal activity events of large and small sizes are highlighted with yellow boxes.

**Figure S3.**
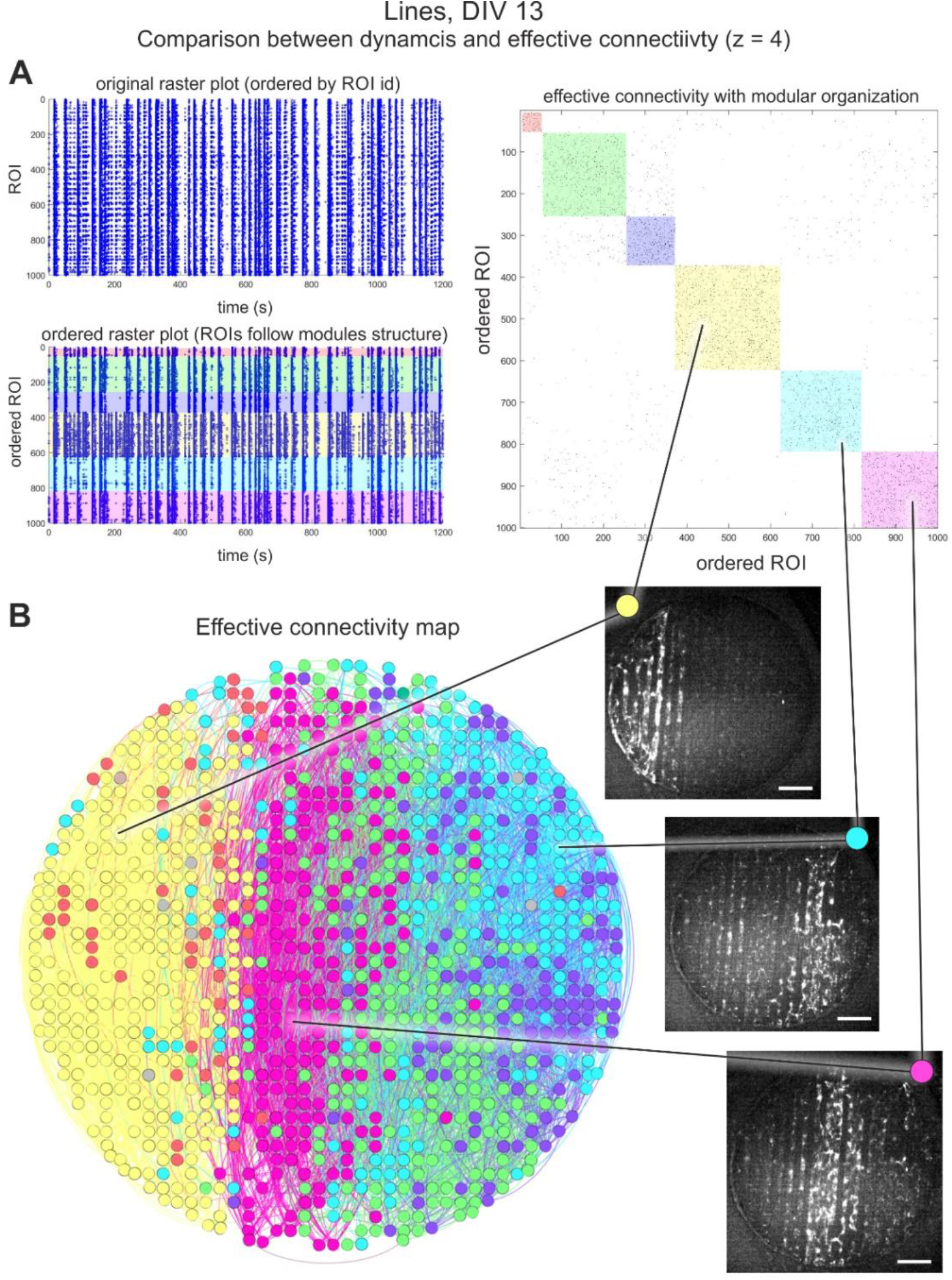
Relation between effective connectivity and dynamics in the ‘tracks’ configuration, and for a significance threshold z=4 to highlight activity outside whole network bursts. (Related to Figure 5)

(A) Original raster plot of spontaneous activity (left-top) and effective connectivity matrix depicting the functional modules (right). The reordering of ROI indexes according to the modules shapes a new raster plot (left-bottom) in which dynamic features of the network can be linked to characteristic modules. The yellow module for instance corresponds to a group of ROIs (reordered indexes 400 to 600) with strong spontaneous activity and the presence of collective events that do not exist in the other groups. This indicates that functional modules reflect specific activity patterns. (B) Network map of effective connections. ROIs are colored according to the functional module they belong to. Some modules, particularly the yellow and the pink, coincide in spatial structure with activity patterns that appear often in the recording (fluorescence snapshots), which illustrates the link between activity and functional organization.

## Notes

### Competing Interest Statement

The authors have declared no competing interest.

## REFERENCES

Aebersold, M.J., Dermutz, H., Forró, C., Weydert, S., Thompson-Steckel, G., Vörös, J., Demkó, L., 2016. “Brains on a chip”: Towards engineered neural networks. TrAC - Trends Anal. Chem. 78, 60–69. https://doi.org/10.1016/j.trac.2016.01.025

Basso, J.M.V., Yurchenko, I., Simon, M., Rizzo, D.J., Staii, C., 2019. Role of geometrical cues in neuronal growth. Phys. Rev. E 99, 1–12. https://doi.org/10.1103/PhysRevE.99.022408

Bisio, M., Bosca, A., Pasquale, V., Berdondini, L., Chiappalone, M., 2014. Emergence of bursting activity in connected neuronal sub-populations. PLoS One 9, e107400. https://doi.org/10.1371/journal.pone.0107400

Blankenship, A.G., Feller, M.B., 2010. Mechanisms underlying spontaneous patterned activity in developing neural circuits. Nat. Rev. Neurosci. 11, 18–29. https://doi.org/10.1038/nrn2759

Bonifazi, P., Difato, F., Massobrio, P., Breschi, G.L., Pasquale, V., Levi, T., Goldin, M., Bornat, Y., Tedesco, M., Bisio, M., Kanner, S., Galron, R., Tessadori, J., Taverna, S., Chiappalone, M., 2013. In vitro large-scale experimental and theoretical studies for the realization of bi-directional brain-prostheses. Front. Neural Circuits 7, 1–19. https://doi.org/10.3389/fncir.2013.00040

Bressloff, P.C., 2000. Traveling waves and pulses in a one-dimensional network of excitable integrate-and-fire neurons. J. Math. Biol. 40, 169–98.

Bullmore, E., Sporns, O., 2012. The economy of brain network organization. Nat. Rev. Neurosci. 13, 336–49. https://doi.org/10.1038/nrn3214

Carola, G., Malagarriga, D., Calatayud, C., Pons-Espinal, M., Blasco-Agell, L., Richaud-Patin, Y., Fernandez-Carasa, I., Baruffi, V., Beltramone, S., Molina, E., Dell’Era, P., Toledo-Aral, J.J., Tolosa, E., Muotri, A.R., Garcia Ojalvo, J., Soriano, J., Raya, A., Consiglio, A., 2021. Parkinson’s disease patient-specific neuronal networks carrying the LRRK2 G2019S mutation unveil early functional alterations that predate neurodegeneration. npj Park. Dis. 7, 1–14. https://doi.org/10.1038/s41531-021-00198-3

Deco, G., Tononi, G., Boly, M., Kringelbach, M.L., 2015. Rethinking segregation and integration: Contributions of whole-brain modelling. Nat. Rev. Neurosci. 16, 430–439. https://doi.org/10.1038/nrn3963

Downes, J.H., Hammond, M.W., Xydas, D., Spencer, M.C., Becerra, V.M., Warwick, K., Whalley, B.J., Nasuto, S.J., 2012. Emergence of a small-world functional network in cultured neurons. PLoS Comput. Biol. 8, e1002522. https://doi.org/10.1371/journal.pcbi.1002522

Feinerman, O., Segal, M., Moses, E., 2005. Signal propagation along unidimensional neuronal networks. J. Neurophysiol. 94, 3406–16. https://doi.org/10.1152/jn.00264.2005

Finc, K., Bonna, K., He, X., Lydon-Staley, D.M., Kühn, S., Duch, W., Bassett, D.S., 2020. Dynamic reconfiguration of functional brain networks during working memory training. Nat. Commun. 11, 1–15. https://doi.org/10.1038/s41467-020-15631-z

Forró, C., Thompson-Steckel, G., Weaver, S., Weydert, S., Ihle, S., Dermutz, H., Aebersold, M.J., Pilz, R., Demkó, L., Vörös, J., 2018. Modular microstructure design to build neuronal networks of defined functional connectivity. Biosens. Bioelectron. 122, 75–87. https://doi.org/10.1016/j.bios.2018.08.075

Golomb, D., Ermentrout, G.B., 1999. Continuous and lurching traveling pulses in neuronal networks with delay and spatially decaying connectivity. Proc. Natl. Acad. Sci. U. S. A. 96, 13480–13485. https://doi.org/10.1073/pnas.96.23.13480

Holloway, P.M., Willaime-Morawek, S., Siow, R., Barber, M., Owens, R.M., Sharma, A.D., Rowan, W., Hill, E., Zagnoni, M., 2021. Advances in microfluidic in vitro systems for neurological disease modeling. J. Neurosci. Res. 99, 1276–1307. https://doi.org/10.1002/jnr.24794

Isaacson, J.S., Scanziani, M., 2011. How inhibition shapes cortical activity. Neuron 72, 231–43. https://doi.org/10.1016/j.neuron.2011.09.027

Jacobi, S., Moses, E., 2007. Variability and corresponding amplitude-velocity relation of activity propagating in one-dimensional neural cultures. J. Neurophysiol. 97, 3597–3606. https://doi.org/10.1152/jn.00608.2006

Jacobi, S., Soriano, J., Moses, E., 2010. BDNF and NT-3 increase velocity of activity front propagation in unidimensional hippocampal cultures. J. Neurophysiol. 104, 2932–9. https://doi.org/10.1152/jn.00002.2010

Kirwan, P., Turner-Bridger, B., Peter, M., Momoh, A., Arambepola, D., Robinson, H.P.C., Livesey, F.J., 2015. Development and function of human cerebral cortex neural networks from pluripotent stem cells in vitro. Development 142, 3178–3187. https://doi.org/10.1242/dev.123851

Li, S., Kuddannaya, S., Chuah, Y.J., Bao, J., Zhang, Y., Wang, D., 2017. Combined effects of multi-scale topographical cues on stable cell sheet formation and differentiation of mesenchymal stem cells. Biomater. Sci. 5, 2056–2067. https://doi.org/10.1039/c7bm00134g

Li, W., Xu, Z., Huang, J., Lin, X., Luo, R., Chen, C.-H., Shi, P., 2014. NeuroArray: a universal interface for patterning and interrogating neural circuitry with single cell resolution. Sci. Rep. 4, 4784. https://doi.org/10.1038/srep04784

Liu, W., Fu, W., Sun, M., Han, K., Hu, R., Liu, D., Wang, J., 2021. Straightforward neuron micropatterning and neuronal network construction on cell-repellent polydimethylsiloxane using microfluidics-guided functionalized Pluronic modification. Analyst 146, 454–462. https://doi.org/10.1039/d0an02139c

Marconi, E., Nieus, T., Maccione, A., Valente, P., Simi, A., Messa, M., Dante, S., Baldelli, P., Berdondini, L., Benfenati, F., 2012. Emergent functional properties of neuronal networks with controlled topology. PLoS One 7, e34648. https://doi.org/10.1371/journal.pone.0034648

Meunier, D., Lambiotte, R., Bullmore, E.T., 2010. Modular and hierarchically modular organization of brain networks. Front. Neurosci. 4, 200. https://doi.org/10.3389/fnins.2010.00200

Millet, L.J., Gillette, M.U., 2012. New perspectives on neuronal development via microfluidic environments. Trends Neurosci. 35, 752–61. https://doi.org/10.1016/j.tins.2012.09.001

Neto, E., Leitão, L., Sousa, D.M., Alves, C.J., Alencastre, I.S., Aguiar, P., Lamghari, M., 2016. Compartmentalized microfluidic platforms: The unrivaled breakthrough of in vitro tools for neurobiological research. J. Neurosci. 36, 11573–11584. https://doi.org/10.1523/JNEUROSCI.1748-16.2016

Nikolakopoulou, P., Rauti, R., Voulgaris, D., Shlomy, I., Maoz, B.M., Herland, A., 2021. Recent progress in translational engineered in vitro models of the central nervous system. Brain 143, 3181–3213. https://doi.org/10.1093/BRAIN/AWAA268

Okujeni, S., Egert, U., 2019. Self-organization of modular network architecture by activity-dependent neuronal migration and outgrowth. Elife 8, 1–29. https://doi.org/10.7554/eLife.47996

Okujeni, S., Kandler, S., Egert, U., 2017. Mesoscale architecture shapes initiation and richness of spontaneous network activity. J. Neurosci. 2552–16. https://doi.org/10.1523/JNEUROSCI.2552-16.2017

Orlandi, J.G., Soriano, J., Alvarez-Lacalle, E., Teller, S., Casademunt, J., 2013. Noise focusing and the emergence of coherent activity in neuronal cultures. Nat. Phys. 9, 582–590. https://doi.org/10.1038/nphys2686

Park, H.J., Friston, K., 2013. Structural and functional brain networks: From connections to cognition. Science (80-.). 342. https://doi.org/10.1126/science.1238411

Park, M.U., Bae, Y., Lee, K.S., Song, J.H., Lee, S.M., Yoo, K.H., 2021. Collective dynamics of neuronal activities in various modular networks. Lab Chip 21, 951–961. https://doi.org/10.1039/d0lc01106a

Pas, S.P., 2018. The rise of three-dimensional human brain cultures. Nature. https://doi.org/10.1038/nature25032

Poli, D., Pastore, V.P., Massobrio, P., 2015. Functional connectivity in in vitro neuronal assemblies. Front. Neural Circuits 9. https://doi.org/10.3389/fncir.2015.00057

Schroeter, M.S., Charlesworth, P., Kitzbichler, M.G., Paulsen, O., Bullmore, E.T., 2015. Emergence of Rich-Club Topology and Coordinated Dynamics in Development of Hippocampal Functional Networks In Vitro. J. Neurosci. 35, 5459–5470. https://doi.org/10.1523/JNEUROSCI.4259-14.2015

Shein-Idelson, M., Ben-Jacob, E., Hanein, Y., 2011. Engineered neuronal circuits: a new platform for studying the role of modular topology. Front. Neuroeng. 4, 10. https://doi.org/10.3389/fneng.2011.00010

Sporns, O., 2013. Network attributes for segregation and integration in the human brain. Curr. Opin. Neurobiol. 23, 162–171. https://doi.org/10.1016/j.conb.2012.11.015

Stiso, J., Bassett, D.S., 2018. Spatial Embedding Imposes Constraints on Neuronal Network Architectures. Trends Cogn. Sci. 22, 1127–1142. https://doi.org/10.1016/j.tics.2018.09.007

Suárez, L.E., Markello, R.D., Betzel, R.F., Misic, B., 2020. Linking Structure and Function in Macroscale Brain Networks. Trends Cogn. Sci. 24, 302–315. https://doi.org/10.1016/j.tics.2020.01.008

Sukenik, N., Vinogradov, O., Weinreb, E., Segal, M., Levina, A., 2021. Neuronal circuits overcome imbalance in excitation and inhibition by adjusting connection numbers. https://doi.org/10.1073/pnas.2018459118

Sumi, T., Yamamoto, H., Hirano-Iwata, A., 2020. Suppression of hypersynchronous network activity in cultured cortical neurons using an ultrasoft silicone scaffold. Soft Matter 16, 3195–3202. https://doi.org/10.1039/c9sm02432h

Tibau, E., Ludl, A.A., Rudiger, S., Orlandi, J.G., Soriano, J., 2020. Neuronal Spatial Arrangement Shapes Effective Connectivity Traits of in vitro Cortical Networks. IEEE Trans. Netw. Sci. Eng. 7, 435–448. https://doi.org/10.1109/TNSE.2018.2862919

Wheeler, B.C., Brewer, G.J., 2010. Designing Neural Networks in Culture: Experiments are described for controlled growth, of nerve cells taken from rats, in predesigned geometrical patterns on laboratory culture dishes. Proc. IEEE. Inst. Electr. Electron. Eng. 98, 398–406. https://doi.org/10.1109/JPROC.2009.2039029

Yamamoto, H., Moriya, S., Ide, K., Hayakawa, T., Akima, H., Sato, S., Kubota, S., Tanii, T., Niwano, M., Teller, S., Soriano, J., Hirano-Iwata, A., 2018. Impact of modular organization on dynamical richness in cortical networks. Sci. Adv. 4, 1–12. https://doi.org/10.1126/sciadv.aau4914

Zhuang, P., Sun, A.X., An, J., Chua, C.K., Chew, S.Y., 2018. 3D neural tissue models: From spheroids to bioprinting. Biomaterials 154, 113–133. https://doi.org/10.1016/j.biomaterials.2017.10.002

## REFERENCES

Bastian, M., Heymann, S., Jacomy, M., 2009. Gephi: An Open Source Software for Exploring and Manipulating Networks. Proc. Int. AAAI Conf. Web Soc. Media 3, 361–362.

Blondel, V.D., Guillaume, J.L., Lambiotte, R., Lefebvre, E., 2008. Fast unfolding of communities in large networks. J. Stat. Mech. Theory Exp. 2008. https://doi.org/10.1088/1742-5468/2008/10/P10008

Faci-Lázaro, S., Soriano, J., Gómez-Gardeñes, J., 2019. Impact of targeted attack on the spontaneous activity in spatial and biologically-inspired neuronal networks. Chaos 29. https://doi.org/10.1063/1.5099038

Fernández-García, S., Orlandi, J.G., García-Díaz Barriga, G.A., Rodríguez, M.J., Masana, M., Soriano, J., Alberch, J., 2020. Deficits in coordinated neuronal activity and network topology are striatal hallmarks in Huntington’s disease. BMC Biol. 18, 1–16. https://doi.org/10.1186/s12915-020-00794-4

Grewe, B.F., Langer, D., Kasper, H., Kampa, B.M., Helmchen, F., 2010. High-speed in vivo calcium imaging reveals neuronal network activity with near-millisecond precision. Nat. Methods 7, 399–405. https://doi.org/10.1038/nmeth.1453

Latora, V., Marchiori, M., 2001. Efficient behavior of small-world networks. Phys. Rev. Lett. 87, 198701-1-198701–4. https://doi.org/10.1103/PhysRevLett.87.198701

Ludl, A.A., Soriano, J., 2020. Impact of Physical Obstacles on the Structural and Effective Connectivity of in silico Neuronal Circuits. Front. Comput. Neurosci. 14. https://doi.org/10.3389/fncom.2020.00077

Orlandi, J.G., Fernández-García, S., Comella-Bolla, A., Masana, M., García-Díaz Barriga, G., Yaghoobi, M., Canals, J.-M., Colicos, M.A., Davidsen, J., Alberch, J., Soriano, J., 2017. NETCAL: An interactive platform for large-scale, NETwork and population dynamics analysis of CALcium imaging recordings, in: Neuroscience 2017. https://doi.org/doi.org/10.5281/zenodo.1119025

Peixoto, T.P., 2017. Nonparametric Bayesian inference of the microcanonical stochastic block model. Phys. Rev. E 95, 1–21. https://doi.org/10.1103/PhysRevE.95.012317

Peixoto, T.P., 2014. The graph-tool python library, Figshare. DOI:10.6084/m9.figshare.1164194

Rubinov, M., Sporns, O., 2010. Complex network measures of brain connectivity: uses and interpretations. Neuroimage 52, 1059–69. https://doi.org/10.1016/j.neuroimage.2009.10.003

Soriano, J., Rodríguez Martínez, M., Tlusty, T., Moses, E., 2008. Development of input connections in neural cultures. Proc. Natl. Acad. Sci. U. S. A. 105, 13758–63. https://doi.org/10.1073/pnas.0707492105

Stetter, O., Battaglia, D., Soriano, J., Geisel, T., 2012. Model-free reconstruction of excitatory neuronal connectivity from calcium imaging signals. PLoS Comput. Biol. 8, e1002653. https://doi.org/10.1371/journal.pcbi.1002653

Tibau, E., Valencia, M., Soriano, J., 2013. Identification of neuronal network properties from the spectral analysis of calcium imaging signals in neuronal cultures. Front. Neural Circuits 7, 199. https://doi.org/10.3389/fncir.2013.00199

